# Natural variants with 2D correlation genetics identify domains coordinating sarcomere proteins during contraction

**DOI:** 10.1101/2021.03.14.435341

**Authors:** Thomas P. Burghardt

## Abstract

Muscle proteins assemble in a sarcomere then by coordinated action produce contraction force to shorten muscle. In the human heart ventriculum, cardiac myosin motor (βmys) repetitively converts ATP free energy into work. Cardiac myosin binding protein C (MYBPC3) in complex with βmys regulates contraction power generation. Their bimolecular complex βmys/MYBPC3 models the contractile system and is used here to study protein coupling. The database for single nucleotide variants (SNVs) in βmys and MYBPC3 surveys human populations worldwide. It consistently records SNV physical characteristics including substituted residue location in the protein functional domain, the side chain substitution, substitution frequency, and human population group, but inconsistently records SNV implicated phenotype and pathology outcomes. A selected consistent subset of the data trains and validates a feed-forward neural network modeling the contraction mechanism. The full database is completed using the model then interpreted probabilistically with a discrete Bayes network to give the SNV probability for a functional domain location given pathogenicity and human population. Co-domains, intra-protein domains coupling βmys and MYBPC3, are identified by their population correlated SNV probability product for given pathogenicity. Divergent genetics in human populations identify co-domain correlates in this method called 2D correlation genetics. Pathogenic and benign SNV data identify three critical regulatory sites, two in MYBPC3 with links to several domains across the βmys motor, and, one in βmys with links to the known MYBPC3 regulatory domain. Critical sites in MYBPC3 are hinges (one known another proposed) sterically enabling regulatory interactions with βmys. The critical site in βmys is the actin binding C-loop, a contact sensor triggering actin-activated myosin ATPase and contraction velocity modulator coordinating also with actin bound tropomyosin. C-loop and MYBPC3 regulatory domain linkage potentially impacts multiple functions across the contractile system. Identification of co-domains in a binary protein complex implies a capacity to estimate spatial proximity constraints for specific dynamic protein interactions in vivo opening another avenue for protein complex structure/function determination.

---

Muscle proteins in the sarcomere assemble to form machinery needed to generate and regulate muscle contraction by their coordinated action. In the human heart ventriculum, cardiac ventricular myosin (βmys) is the molecular motor repetitively converting ATP free energy into work. Three genes, MYH7 (heavy chain), MYL3 (essential light chain or ELC), and MYL2 (regulatory light chain or RLC) encode βmys (**Fig 1**). The heavy chain has a 140 kDa N-terminal globular motor domain (subfragment 1 or SF1) and an extended α-helical tail domain forming subfragment 2 (S2) and light meromyosin (LM). LM domains form dimers and self-assemble into myosin thick filaments with SF1 and S2 projecting outward from the core in a helical array ^*1*^. Thick filaments interdigitate with actin thin filaments (filamentous or F-actin) in the sarcomere where βmys motors cyclically interact with F-actin ^*2*^. SF1 binds to an actin filament and rotates the lever arm generating torque and tension on F-actin ^*3, 4*^. The ELC N-terminus also binds actin generating force ending the power stroke. Fixed myosin generates tension translating actin towards the pointed end and against the loading force (**Fig 1**).

**Figure 1.**
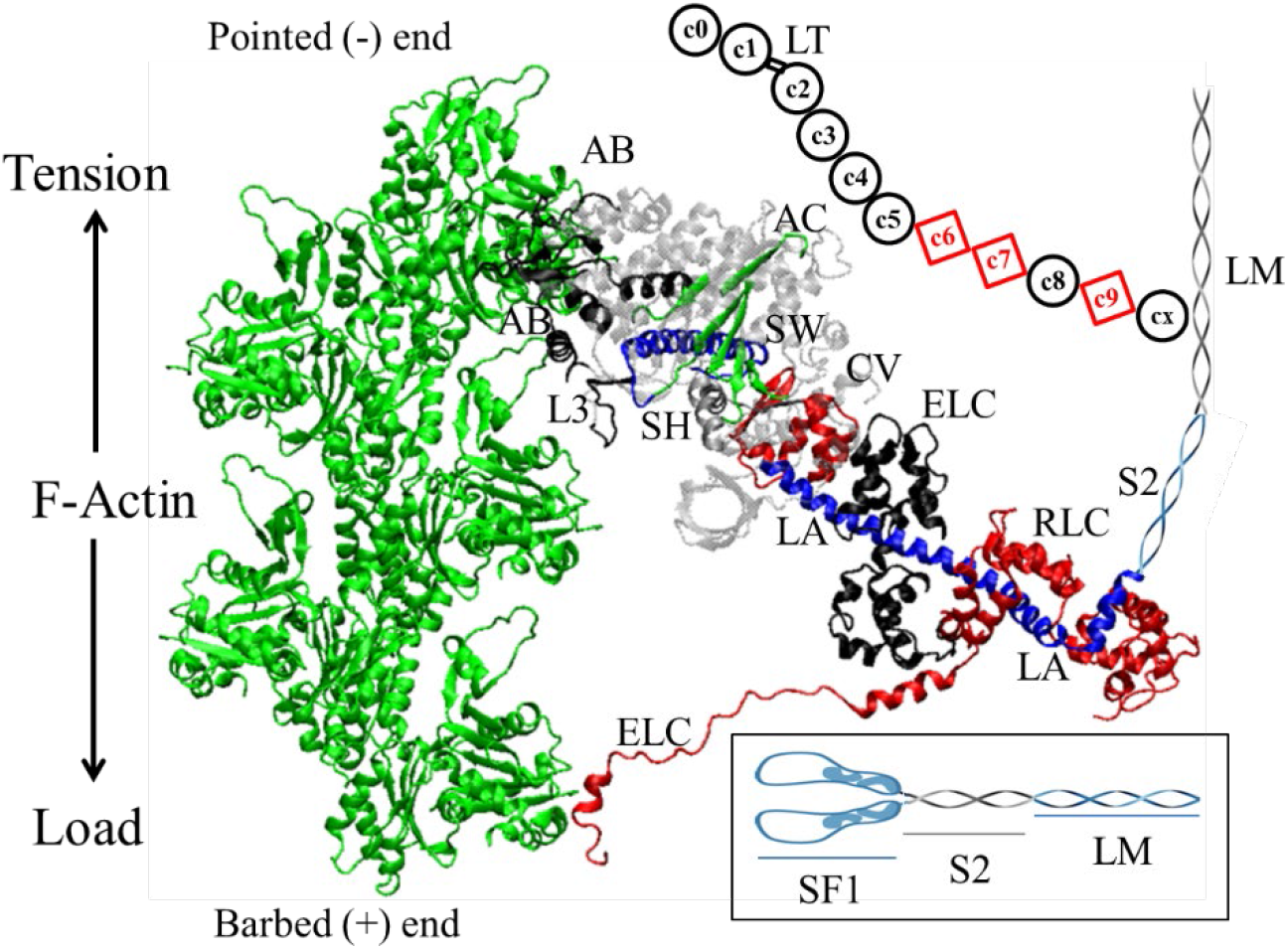
Myosin dimer (insert) has two subfragment 1 (SF1), subfragment 2 (S2), and light meromyosin (LM) polypeptides. SF1 has a motor domain and a lever arm (LA, blue) with bound light chains ELC (black) and RLC (red). MYBPC3 (not drawn to scale) has multiple domains shown in black circles or red squares indicating immunoglobulin-like or fibronectin-like domains with CX binding myosin LM and N-terminal C0-C2 maintaining transient interactions with actin (green) and myosin.

The lever arm complex, stabilized by bound essential and regulatory light chains (ELC and RLC) ^*5, 6, 7*^, converts torque generated by ATP hydrolysis in the motor into linear displacement of myosin vs actin filaments. The relative movement of motor subdomains is the structural manifestation of the energy conversion wherein ATP hydrolysis at the active site (AC, **Fig 1**) modulates actin affinity at the actin binding sites (including CL, ML, L2, L3, and AB, **Figs 1 & 2**), induces small active site translations amplified into larger structural changes at the switch 2 helix (SW) and SH1/SH2 hinge (SH), and the latter converted by the converter domain (CV) into the large lever arm (LA) swing.

**Figure 2.**
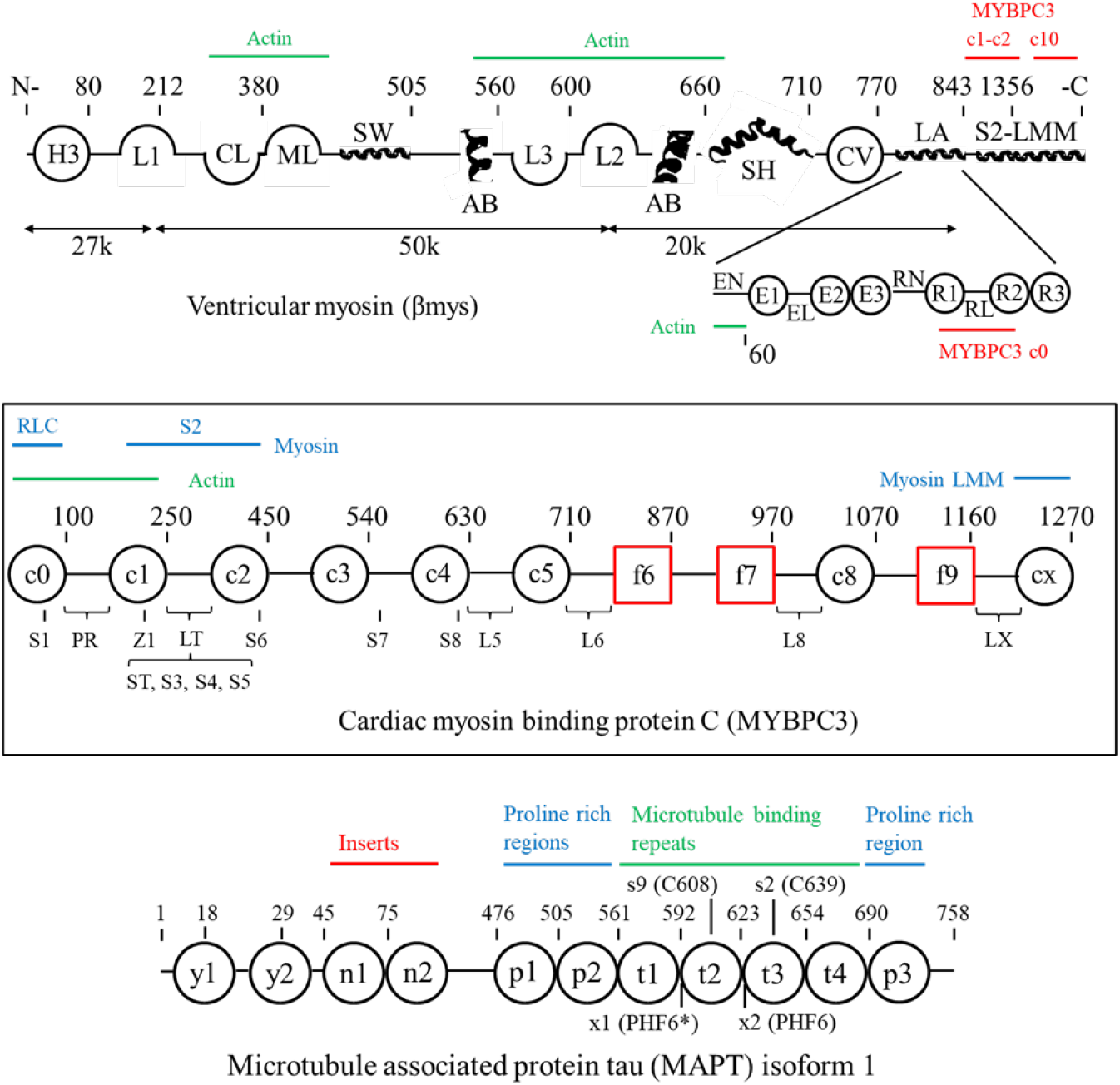
Linearized diagrams for the βmys, MYBPC3, and MAPT structures identifying most domains defined in **SI Tables S1 & S4. Top**. The βmys diagram does not indicate the active site (AC), OM binding site (OM), and mesa (ME) because they occupy multiple regions in the linearized representation. Myosin light chains appear below the heavy chain. MYBPC3 and actin binding sites are indicated in red and green above the heavy chain and below the light chains. **Middle**. The MYBPC3 diagram has 8 Ig-like domains (black circles) and 3 fibronectin-like domains (red squares). Serine phosphorylation sites, S1 (Ser47), ST (Ser273), S3 (Ser282), S4 (Ser302), and S5-S7 and a threonine phosphorylation site (S8) are indicated below the chain. Domain linkers of interest include the proline rich linker (PR) and LT containing a regulatory site. Z1 is a zinc binding site. Myosin S2 and LM and actin binding sites on MYBPC3 are indicated above the linearized model. **Bottom**. The MAPT diagram is for the largest isoform (isoform 1). Sites or domains identified are tyrosine phosphorylation sites (Y1 and Y2), N-terminal inserts (N1 and N2), proline rich regions (P1-P3), microtubule binding repeats (T1-T4), cysteines C608 and C639 (S9 and S2), and hexapeptide motifs PHF6* (VQIINK at x1) and PHF6 (VQIVYK at x2).

Cardiac myosin binding protein C (MYBPC3), denoting either the gene or expressed protein depending on context, localizes to the C-zone in the cardiac muscle sarcomere where it regulates energy conversion and shortening velocity by transient N-terminus binding to actin or myosin with the C-terminus fixed to the myosin thick filament ^*8*^. MYBPC3 has a linear array of 8 immunoglobulin-like (Ig-like) and 3 fibronectin-like domains (**Fig 2**). Peptide linker 2 (LT), in the N-terminus linking Ig-like domains C1 and C2, contains phosphorylation sites participating in contractile regulation by modulating myosin activity ^*9-11*^ and calcium sensitivity ^*12*^. The MYBPC3 N-terminus associates alternatively with thin filament actin ^*13, 14*^, the myosin motor ^*15*^, and myosin thick filament ^*13*^ while the C-terminal domain binds to LM in the thick filament with CX ^*16*^. Two hinges in the linear molecule, at LT and C5, were identified ^*17*^. A plausible third hinge near the C-terminus would facilitate the known multiple protein interactions with βmys. βmys, MYBPC3, and actin form the elements defining muscle function in the heart ^*18*^. Cardiac myosin binding protein C (MYBPC3) in complex with βmys regulates contraction power generation. Their bimolecular complex βmys/MYBPC3 is a model for the contractile system and for studying interactions coupling them.

Non-synonymous single nucleotide variations (SNVs) change protein sequences in βmys and MYBPC3. Sequence variants modify protein domain structure and function affecting phenotype and pathogenicity outcomes. Moreover, βmys and MYBPC3 SNVs have pathogenicities that correlate with human populations ^*19*^ indicating that cardiac muscle physiology integrates βmys/MYBPC3 complex functionality with the wider system involving genetic background ^*20*^. It is proposed that genetic background influence can be leveraged to identify intra-protein coordinated functional domains (co-domains) that are responsible for coordinating the βmys/MYBPC3 complex function.

The database for SNVs in the βmys and MYBPC3 consistently records SNV physical characteristics including substituted residue location in the protein functional domain, the side chain substitution, substitution frequency, and human population group, but inconsistently identifies SNV implicated phenotype and pathology outcomes. A selected consistent subset of the data trains and validates a feed-forward neural network modeling the contraction mechanism. The database is completed using the model then interpreted probabilistically with a discrete Bayes network to give the SNV probability for a functional domain location given pathogenicity and human population as described earlier ^*19, 21*^. Co-domains, intra-protein domains coupling βmys and MYBPC3, are identified by their population correlated SNV probability product for given pathogenicity. Divergent genetics in human populations identify co-domain correlates in this method called 2D correlation genetics.

Human population genetic divergence decreases linearly with increasing human migration distance over the earth’s surface from a single origin in Africa ^*22*^. Migration distance is then a proxy for genetic differentiation providing the means for population indexing and 2D correlation genetics mapping intra-protein co-domain correlations and by extension co-domain interactions. 2D correlation genetics is an approach analogous to 2D correlation spectroscopy ^*23*^ wherein βmys and MYBPC3 functional domains replace the two spectral frequencies near resonance, SNV probability correlates mimics resonance absorption intensity, and human population genetic differentiation provides the arithmetically indexed perturbation.

2D NMR uses the cross-correlated response of specific nuclei to the perturbation provided by selected radio frequency (RF) pulse sequences and magnetization transfer for the purpose of molecular structure determination. In concept it is analogous to 2D correlation genetics with the latter supplying qualitative proximity constraints. Identification of co-domains in a multiprotein complex implies a potential to use proximity constraints for elaborating dynamic protein interactions in vivo via structural ensembles provided by constrained molecular dynamics simulation also analogous to 2D-NMR based structure determination. 2D correlation genetics maps path-of-influence mechanical coupling across the βmys/MYBPC3 complex ^*24*^. These ideas and findings define a new genetic approach for locating bilateral structure/function relationships in a multiprotein complex.

## 2. METHODS

### 2.1 SNV data retrieval

Data retrieval is described for the myosin heavy chain gene MYH7. Protocols for other genes involved including MYL2, MYL3, MYBPC3, and MAPT (microtubule associated protein tau) are identical after substituting the appropriate gene name. Using a browser, open the National Center for Bioinformatics (NCBI) home page, select the SNP database, and use “MYH7 missense_variant” as the search item. The search returns multiple items saved to a text file that is subsequently imported to the Mathematica (Wolfram, Champaign, IL, USA) program Sort. Sort uses text search/extract tools to identify and collect the SNV reference numbers (rs#) into a list. Redundant rs# entries are merged by the program into one rs#. Each rs# corresponds to a single reference gene location and nucleotide but may involve various nucleotide substitutions. Sort uses rs#’s in an URL API (eg., https://api.ncbi.nlm.nih.gov/variation/v0/beta/refsnp/2258689) to download JSON format data containing the information normally in view using the browser. JSON data for each rs# is exported to a file.

Exported JSON files are read and interpreted in program SNP to extract data characterizing each SNV in a standardized format. It creates a single matrix containing the information used for training the neural network. JSON data interpretation involves assigning: the sequence number for the substitution, its functional domain in the protein, residue substitution (sidechain), allele frequency, human population, phenotype, and pathology. Unique two letter codes define each domain, side chain substitution, phenotype, and pathology. Unique three letter codes define each human population.

### 2.2 Neural/Bayes network configuration

Trial network configurations associate mutant (mu) location in the protein domain (cd), residue substitution (re), population group (po), and SNP allele frequency (af) in a causal relationship with phenotype (ph) and pathogenicity (pa) in a directed acyclic graph (DAG) in **Fig 3 panel a** and as described previously ^*19*^.

**Figure 3.**
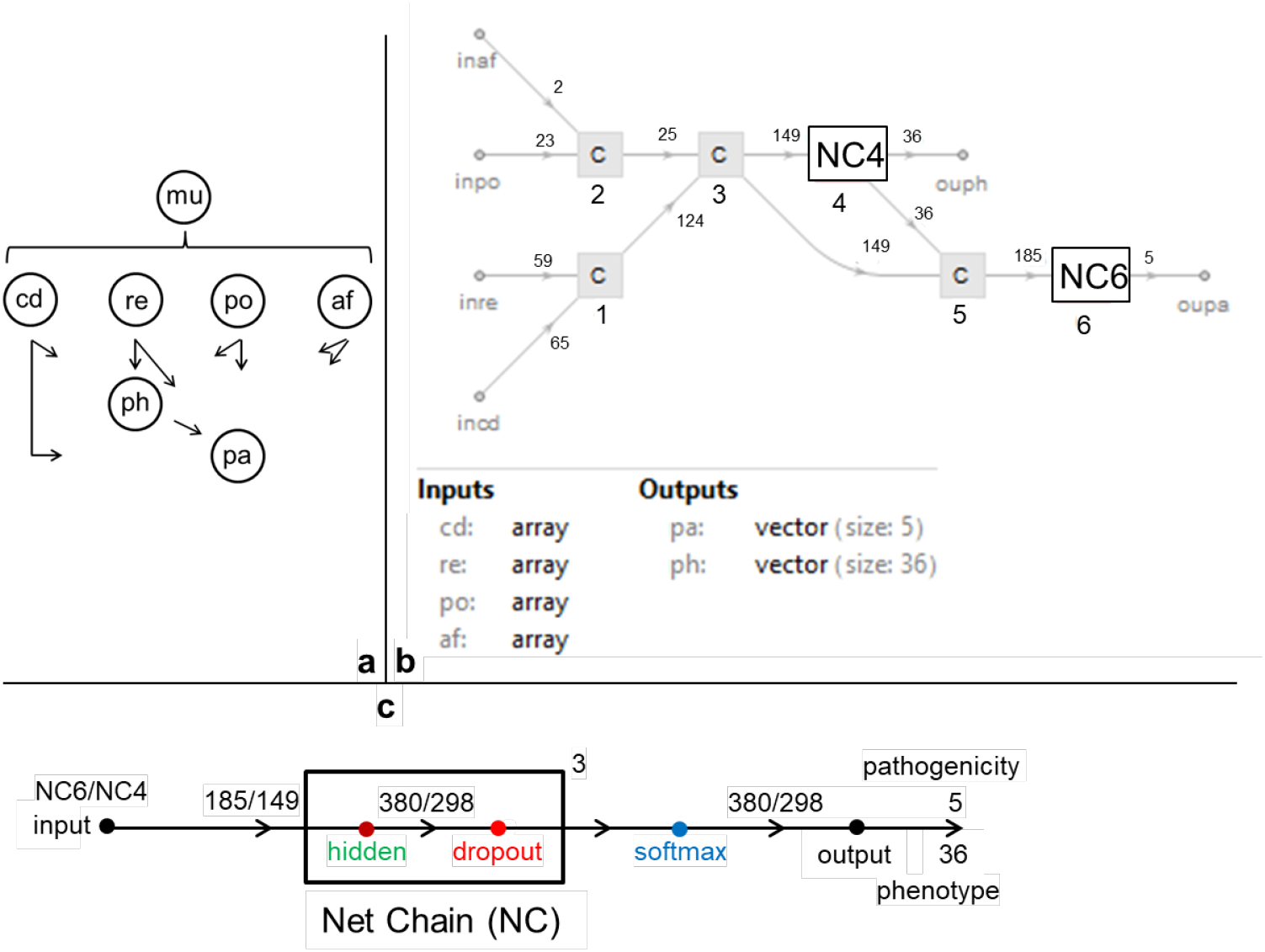
The neural network modeling structure/function influences from heart disease. **Panel a**. Directed acyclic graph (DAG) denoting relationships among mutant (mu) domain location (cd), residue substitution (re), human population (po), allele frequency (af), phenotype (ph), and pathogenicity (pa) modeling the contraction mechanism with neural and Bayes networks. **Panel b**. Feed forward neural net corresponding to the directed acyclic graph (DAG) model for contraction (from **panel a**) and relating inputs (prefix in) for the site of the SNV modification (cd), residue substitution (re), human population (po), and allele frequency (af) with outputs (prefix ou) for disease phenotype (ph) and pathogenicity (pa). Components denoted by C (1-3, and 5) catenates input lists to Net Chains denoted with NC (components 4 and 6) defined in **panel c**. Numbers near arrows in the lines input to or output from C and NC components are the number of nodes. Node number shown are for βmys/MYBPC3 in complex with 65 domains and 59 ref/sub pairs input to C1, and, 23 human population categories and 2 allele frequency categories input to C2. Output pathogenicities always number 5. **Panel c**. Net Chains (NC) are three connected hidden/dropout layers (superscripted 3) that output to a softmax layer described in the text. Numbers above the arrows in the horizontal line are nodes for NC6/NC4 in **panel b**. Node numbers are for the βmys/MYBPC3 complex.

Protein domains (cd in **Fig 3 panel a**) and their 2 letter abbreviations are indicated in Supplementary Information (**SI**) **Table S1**. A protein complex made from four genes, MYH7, MYL2, MYL3, and MYBPC3 has domains (cd) from 65 functional sites combining assignments made previously ^*21*^ and new assignments in myosin including blocked head/converter interface (BH, index 3) ^*25*^, mesa trail (MR, index 17) ^*26*^, binding sites for C1, C2, and LT in MYBPC3 on myosin S2 (M1, M3, and M2, indices 18, 20, and 19) ^*27, 28*^, for CX in MYBPC3 on myosin LM (M5, index 21) ^*29*^, binding sites for titin and myomesin on myosin LM (T1 and Y1, indices 28 and 29) ^*29-31*^, and for subdomains in RLC (RN, R1, RL, R2, and R3, indices 35-39) ^*32*^. Every SNP in the database has an assigned domain. **Fig 2** shows linear representations of myosin and MYBPC3 indicating mutual binding sites and the locations of most domains listed in **SI Table S1**.

Residue substitution (re in **Fig 3 panel a**) refers to the SNV reference and substituted residue (ref/sub) pair. Ref/sub combinations have 420 possibilities for 21 amino acids. Combinations reduced to 59 when side chains are pooled into categories related to size, hydrophilicity, and charge such that (Arg, His) ⟶Lys, Asp ⟶Glu, Ser ⟶Thr, Asn ⟶Gln, (Ala, Val, Ile, Met, Phe, Trp) ⟶Leu.

Human population group (po in **Fig 3 panel a**) and residue substitution prevalence (allele frequency or af in **Fig 3 panel a**) fill out the independent parameters in the network. Populations and their 3 letter abbreviations are indicated in **SI Table S2**. Population categories in **SI Table S2** are self-evident or are defined in the caption. Allele frequency is a continuous variable in the database on the interval 0 ≤ af ≤ 1 for 1 meaning all alleles are substituted by the SNV. These data are divided into two discrete categories of ≤1% (category 0) or >1% (category 1).

Phenotype (ph) data classifications for cardiovascular disease pertaining to βmys and MYBPC3 variants has 36 categories in the NCBI SNP database. Names and two letter codes are listed in **SI Table S3**. All but 2 phenotypes associate with both βmys and MYBPC3 SNV’s. The two exceptions are noted in parenthesis. NCBI phenotype categories have changed significantly over time and occasionally conflict for a given SNV for data submissions from different sources. One phenotype from the pool is assigned when conflicting selections have a clearly dominant choice. In all other cases the unknown category is assigned.

Pathogenicity (pa) data have standardized classifications in the NCBI SNP database that includes pathogenic (pt), likely pathogenic (lp), benign (be), likely benign (lb), and unknown (uk). The pathogenicities and their 2 letter codes are summarized in **Table 1**. NCBI pathogenicity category assignment from different sources sometimes conflict. One pathogenicity is assigned from the pool when there is a clearly dominant choice. In all other cases the unknown category is assigned.

**Table 1.**
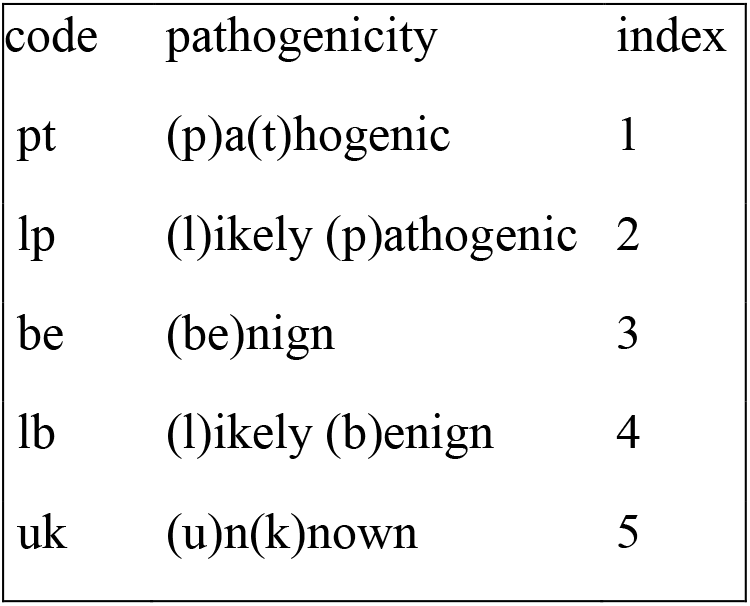
Pathogenicity (pa) with 2 letter codes.

| code | pathogenicity | index |
| --- | --- | --- |
| pt | (p)a(t)hogenic | 1 |
| lp | (l)ikely (p)athogenic | 2 |
| be | (be)nign | 3 |
| lb | (l)ikely (b)enign | 4 |
| uk | (u)n(k)nown | 5 |

### 2.3 Neural network validation

The neural network indicated in **Fig 3 panels b & c** models structure/function influences from disease and follows from the DAG in **Fig 3 panel a**. The neural net relates domain position (cd), residue substitution (re), population (po), and allele frequency (af) inputs to phenotype (ph) and pathogenicity (pa) outputs through 3 fully connected linear (hidden) and dropout layers with 380 (NC6) or 298 (NC4) nodes each, and, a softmax layer conditioning output for digital classification of phenotype and pathogenicity. Dropout layers mitigate overtraining.

The NCBI SNP database was mined to assign the known independent and dependent discrete variables for βmys and MYBPC3 in 6-dimensional data points (fulfilled 6ddps). The latter train and validate the neural network that can then predict phenotype and pathogenicity for any single residue substitution in complexed βmys/MYBPC3. The SNP database also contains a majority of 6ddps having one or both dependent data points unknown (unfulfilled 6ddps). They are predicted by the neural network model. Independent discrete variables are always knowns in the 6ddps.

All fulfilled 6ddps are validating-data. Training-data contains 49.5% of the validating-data, complement-data contains 50% of the validating-data not including training-data, and pseudo-new-data is the remaining 0.5% of validating-data not in training-or complement-data. Trial training-data sets of fulfilled 6ddps are selected randomly but subject to the constraint that each phenotype and pathogenicity outcome in the fulfilled 6ddps is represented in the training set except when their total representation in the fulfilled 6ddp’s is <2 occurrences. Training-data is chosen first. Members of the training data set change with each new trial. Complement-data and pseudo-new-data are then selected randomly from validating-data not in the training set. Complement-data is used to “validate” the model during the optimization phase of training where learnable weights and biases for the linear layers are assigned. Weight initializations are normally distributed with zero mean and standard deviation of (1/n)^½^ (for n inputs). Bias is initialized to zero.

Correctly classifying new SNPs is the only purpose of the optimized neural net model. Model suitability for purpose is tested using the pseudo-new-data. Each pseudo-new-unknown 6ddp has its position and substitution assignment evaluated by the neural network trial with the output phenotype and pathogenicity compared to know values. This comparison is the new-unknown predictor metric used to rank a neural network model for best suitability. The neural network models ranked by their new-unknown predictor metric are best at predicting unfulfilled (ph, pa) outputs from their domain position (cd), residue substitution (re), population (po), and allele frequency (af) assignments.

Best ranked model solutions formed a small subset from the larger pool of independent solutions. Model solution members in the subset were increased (best solutions by the new-unknown predictor metric used first) until collective quantities were unchanged by further enlargement. This selection process favored model subsets sufficiently diverse to cause normally distributed estimates for Bayes network probabilities (discussed below) implying that they randomly sample the set of best implicit transduction models approximating the real transduction mechanism by minimizing random error. A total of ∼2000 trials generate 3-10 best implicit models. This process was repeated until ∼100 total models were collected. These models were combined into 20 best-of-the-best implicit models. These 20 neural networks are distinct implicit models for the role of complex βmys/MYBPC3 in transduction.

The selection process outlined above, suitable for minimizing random error, is unlikely to address systematic model limitations. They exactly reproduce ≥60% of the known 6ddps in the target protein complex constraining potential systematic errors in the models to <40% of the dataset and implying the measure of their accuracy. Systematic error increased substantially together with phenotype category expansion. The difficulty is accounting (with the new new-unknown predictor metric) for a wide variety of assigned phenotypes while just a very few (e.g., hc and cm) predominate (**SI Table S3**). Enlarging the independent parameter set and expanding depth and connectivity in the neural network models are likely to improve model reliability while requiring more computing resources. Here the new approach to investigate intra-protein connectivity in a complex molecular machine is developed. Systematic errors will be addressed later in subsequent work by enlargement of the residue substitution parameter to 420 (vs current 59) possibilities for 21 amino acids and by enlarging depth and connectivity in the neural network models.

Fulfilled and predicted outputs combined are the database for Bayes network (**Fig 3**) tasked with formulating a statistics based complexed βmys/MYBPC3 transduction mechanism as described in the next section.

### 2.4 Bayes network modeling of complexed βmys/MYBPC3 transduction mechanism

**Fig 3 panel a** shows the DAG for the Neural/Bayes network model. Arrows imply a direction for influence hence the domain (cd), residue substitution (re), population (po), and allele frequency (af) assignment implies a probability for phenotype (ph) and pathogenicity (pa). Datasets 6ddpbmysMYBPC3.xls and 6ddpMAPTMYBPC3.xls described in **SI** show the fulfilled and unfulfilled 6ddps for complexed βmys/MYBPC3 and MAPT/MYBPC3 containing 39,290 and 31,181 total variations, respectively. Combined fulfilled and predicted 6ddp data sets define conditional probabilities for the systems in the form of conditional probability tables or CPT’s. The product of conditional probabilities defines the joint probability density,

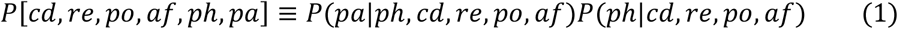

where the right side of eq. 1 is the product of conditional probabilities and the joint probability density on left hand side. Calculating SNV probability for domain *i* in population *j* uses joint probability density,

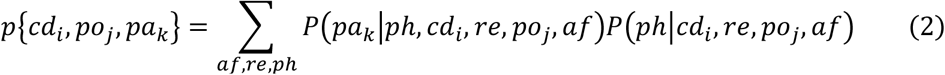

where summation is over all values for allele frequency (af), residue substitution (re), and phenotype (ph). How joint probabilities correlate between domains from two proteins in the complex over population (po) is investigated here with 2-dimensional (2D) correlation testing described in section 2.6.

### 2.5 Bayes network modeling of complex MAPT/MYBPC3

A control computation on a system that presumably never forms a natural functional entity, complex MAPT/MYBPC3, is identical to that done for complexed βmys/MYBPC3. Microtubule-associated protein tau (MAPT) is an intrinsically disordered protein regulating microtubule formation from tubulin ^*33, 34*^. Tau amyloid aggregation associates with tau hyperphosphorylation ^*35*^ and neurodegenerative diseases including Alzheimer’s ^*36*^. Alternative splicing variants of the MAPT gene expresses eight isoforms of the protein (isoforms 1-8) that have SNV’s in the NCBI SNP database. MAPT structural organization for the 758 kDa molecular mass variant (isoform 1) is shown in **Fig 2** identifying tyrosine phosphorylation sites (y1 and y2), N-terminal inserts (N1 and N2), proline rich regions (P1-P3), and microtubule binding repeats (mtbr T1-T4). Tao aggregation beginning with paired helical filament (PHF) formation involves cysteins C608 (S9) and C639 (S2) ^*37*^ and hexapeptide motifs PHF6* (VQIINK at X1) and PHF6 (VQIVYK at X2) ^*38, 39*^. Other isoforms are shorter principally by deletion of t2 with 2, 1, or 0 N-terminal inserts.

Every SNP in the database has an assigned domain (cd). MAPT and MYBPC3 collectively have 42 domains with descriptions and 2 letter codes listed in **SI Table S4**. Population (po) classifications for diseases pertaining to MAPT and MYBPC3 variants have 36 categories in the NCBI SNP database. Names and 3 letter codes are listed in **SI Table S5**. Phenotype (ph) classifications for disease pertaining to MAPT and MYBPC3 variants have 30 categories in the NCBI SNP database. Names and 2 letter codes are listed in **SI Table S6**.

### 2.6 2D correlation testing

SNV’s from human population *po*_*j*_ reside in a protein domain *cd*_*i*_ with probability given by eq. 2. Furthermore, pathogenicity (pa) is made binary, either pathogenic or benign, by combining likely-pathogenic with pathogenic probabilities or likely-benign with

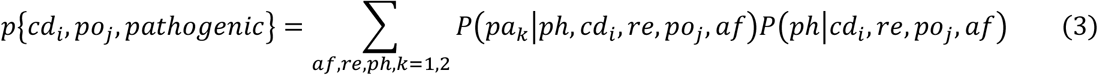

benign probabilities (eq. 2 and k=1,2 or 3,4, respectively, see **Table 1**). In this scenario, eq. 2 specializes into two equations,

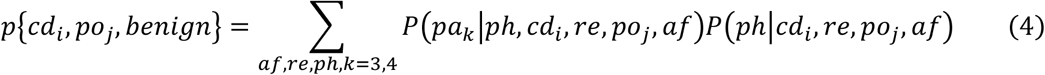

Eqs. 3 & 4 expressions populate the synchronous and asynchronous generalized 2D correlation intensities ^*23*^,

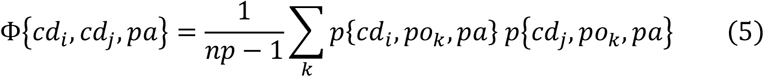

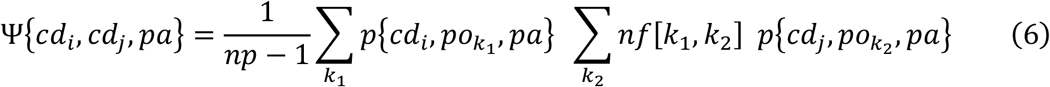

for,

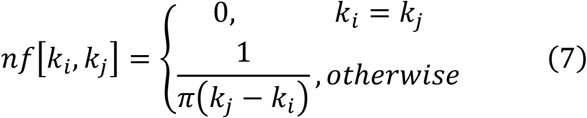

Φ in eq. 5 gives the synchronous, and Ψ in eq. 6 the asynchronous, 2D correlation intensities for *np* the number of populations represented by *po*, and *pa* either pathogenic or benign. The 2D synchronous intensity map will identify domain pairs whose SNV probabilities correlate or anti-correlate for identical populations while the 2D asynchronous intensity map identifies domain pairs whose SNV probabilities lead or lag each other over populations ordered by quantitation of their genetic diversity. The latter calculation is described in Methods section 2.8. Synchronous 2D maps in this application have only intensity correlations (never anti-correlations) implying SNV probability between domain pairs always increase or decrease coincidentally. Consequently, relative phase is not identified for synchronous correlations.

Data obtained from calculation of protein domain SNV probability correlations is represented in 2D contour plots showing Φ or Ψ for the identical listing of first βmys (abbreviated M7) then MYBPC3 (C3) domains, in the order given in **SI Table S1** for a total of 65 domains, on both x-and y-axes. Domains are discrete entities hence the 2D contour plots resemble a pixelated image with grayscale representing intensity. Both inter-and intra-protein correlations are indicated with domain autocorrelated intensity on the diagonal. Cross-protein cross-domain correlations (co-domain correlations), representing C3 vs M7 domains are the focus of this analysis.

The most significant combined synchronous and asynchronous co-domain interaction cross-correlates are the largest elements in the array,

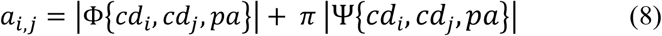

for Φ and Ψ from eqs. 5 and 6, *cd*_*i*_ and *cd*_*j*_ the SNV containing functional domains in βmys and MYBPC3, and *pa* their common pathogenicity (pathogenic or benign). Array element *a*_*i,j*_ is the product of probabilities for SNV’s in each domain member of the co-domain weighted by synchronous or asynchronous coupling to human populations. Quantity π multiplying the asynchronous cross-correlates (Ψ) in eq. 8 balances weighting for synchronous (Φ) with nearest neighbor population asynchronous cross-correlates. Elements *a*_*i*,j_ are ≥ 0. They are combined with *−a*_*i,j*_ then all are plotted in a histogram. Results are a symmetric distribution from 2028 elements well approximated with a single Gaussian distribution giving the standard deviation. The significance of co-domain cross-correlation is measured by its distance from the mean (in this case 0) expressed in multiples of standard deviation. Just the 6 most significant co-domain interactions for complex βmys/MYBPC3 for pathogenic or benign SNVs were interpreted.

Control complex MAPT/MYBPC3 analytics likewise focus on the 6 most significant co-domain interactions. The significance of the findings for each system reflects on the contrast between complexed systems with documented intra-protein contacts (βmys/MYBPC3) compared to one with presumably none (MAPT/MYBPC3).

Population dependence of co-domain interactions in the 6 most significant co-domain interactions is described only for the βmys/MYBPC3 complex using quantities,

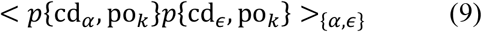

or

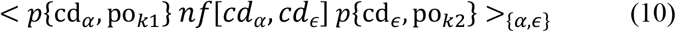

for synchronous (eq. 9) or asynchronous (eq. 10) correlations, < … >_{α,ε}_ implying averaging over 6 most significant co-domains (*cd*_*α*_,*cd*_*ε*_), and population *po*_*k*_. Asynchronous population dependence (eq. 10) involves population pairs (k_1_,k_2_) requiring two dimensions for display.

### 2.7 Co-domain interaction by real or virtual mechanisms

Any co-domain pair from two proteins, A and B, in complex implies there are two functional domains (one in A and one in B) where SNV modification probabilities cross-correlate over genetic diversity in human populations. It occurs by two mechanisms named real and virtual. **Fig 4** shows a hypothetical example for the real mechanism in a complex of A and B where protein A is βmys and protein B is MYBPC3. Consider the functional domains CL and ML in βmys, and, S3 and S7 in MYBPC3. They are SNV modified individually as shown in **panels a-d** (i.e., data in the SNP database contains the SNV frequency data for these species). Favorable cross-correlation of their SNV modification probabilities implies the real co-domain interaction indicated by the solid green lines in **panel f**. SNVs in **panels a-d** might disrupt the real S3-CL or S7-ML co-domain interactions. This is indicated by a red line connecting the functional domains.

**Figure 4.**
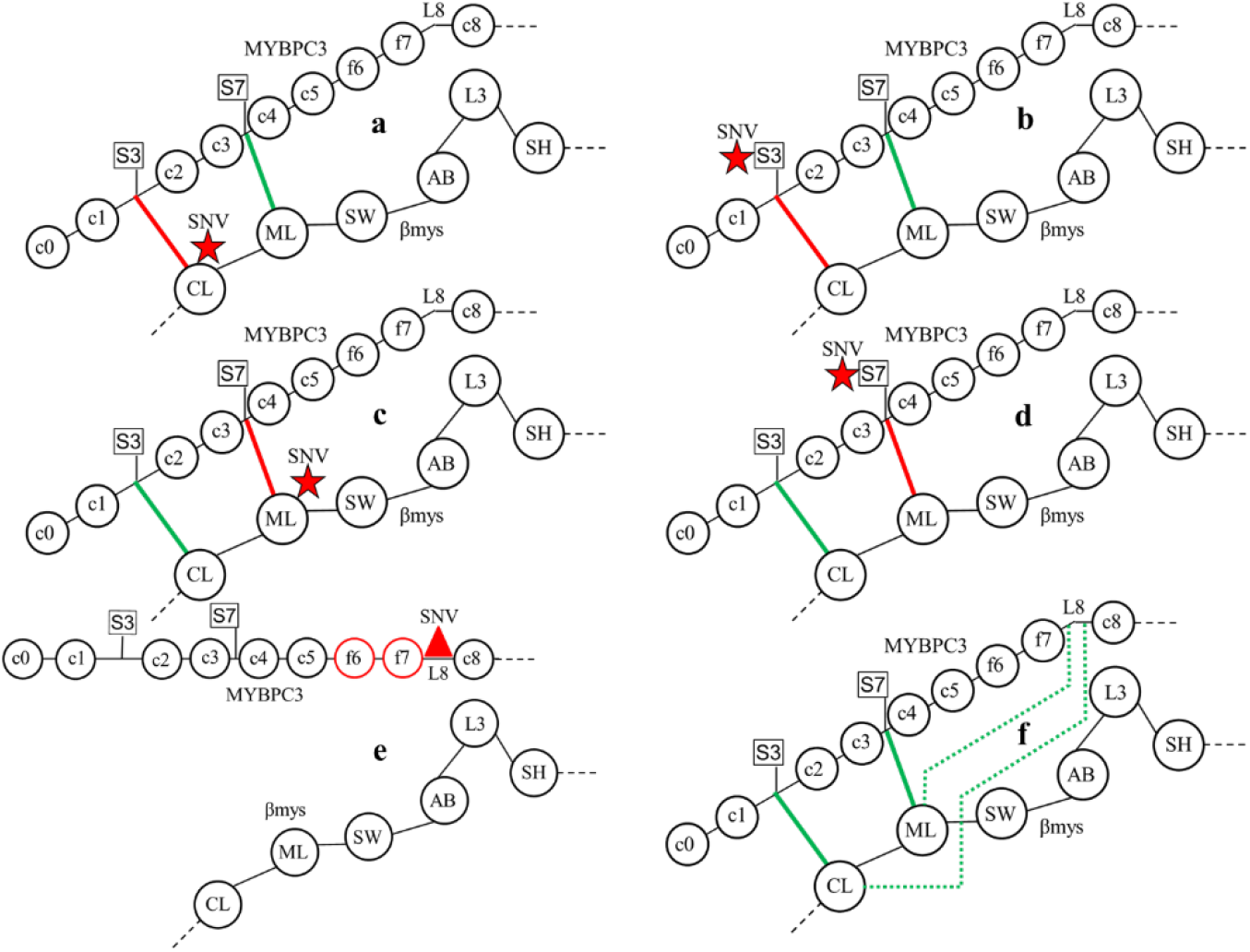
Hypothetical βmys/MYBPC3 complexes in human populations for SNVs detecting real and virtual co-domains. Functional domains CL and ML in βmys, and, functional domains S3 and S7 in MYBPC3 are SNV modified individually as shown in **Panels a-d**. Complexes SNV (star) modified at C-loop (a), phosphorylatable serine S3 (b), myopathy loop (c), or phosphorylatable serine S7 (d) are species needed in the database to surmise the co-domain interactions in the native species shown in Panel f. The native complex involves real co-domains indicated by the green lines between S3 and CL and between S7 and ML. Co-domain interruption by SNVs is indicated by the red lines connecting S3-CL or S7-ML in **Panels a-d**. Real co-domain detection is by cross-correlation of SNV probabilities in species a-d over various human populations. SNV modification in **Panel e** is located in L8 where it alters MYBPC3 conformation to interrupt real co-domains S3-CL and S7-ML. Favorable cross-correlation of the SNV modified species in **panels a, c, and e** implies the virtual co-domains L8-CL and L8-ML indicated by the broken green lines in **panel f**. SNV containing species have identical pathogenicity (benign or pathogenic) but they can represent different phenotypes.

Alternatively, the virtual mechanism has a SNV altered functional domain in protein B that perturbs one or more real co-domain contacts between proteins A and B in complex. The real co-domain contacts do not involve the SNV altered functional domain in protein B. This scenario is also depicted in **Fig 4** for the virtual co-domains in complex βmys/MYBPC3. The SNV modification in **panels e** is located in L8 where it alters MYBPC3 conformation to interrupt real co-domains S3-CL and S7-ML. Favorable cross-correlation of the SNV modified species in **panels a, c**, and **e** implies the virtual co-domains L8-CL and L8-ML indicated by the broken green lines in **panel f**. Inter-protein interactions implied by correlations of SNV modified species in **panels b, d**, and **e** are not interpreted. **Fig 4** shows that detecting the real co-domain combinations, S3-CL and S7-ML, involves detecting the four SNVs at S3, S7, CL and ML. In addition, **Fig 4** shows that detecting the virtual co-domains combinations, L8-CL and L8-ML involves detecting three SNVs at L8, CL, and ML.

Real vs virtual mechanisms for the co-domain interactions identified by SNV probability cross-correlation are indistinguishable without additional information from another source. Examples in **Fig 4** show real co-domains involve a pair of cross-protein functional domains while virtual co-domains involve multiple functional domain interactions. Real co-domains coordinate across complexed proteins requiring minimally a binary cooperation that would confer more specificity, a desirable characteristic. Virtual co-domains lack specificity and readily involve multiple interactions. Independent corroborating structural information for MYBPC3 suggesting C5 is a hinge allowing the linear MYBPC3 molecule to bend adds the context needed to propose the virtual mechanism for co-domains involving C5.

### 2.8 Population genetic diversity proxy

Worldwide human genetic diversity attributed to migration from a single origin in East Africa is based on the serial founder effect addressing migration ^*22*^, colonization, and exchange between geographically near populations ^*40*^. The serial founder effect implies the observed linear diversity decrease with human migration distance over the earth’s surface. Linearly decreasing genetic variation consistent with the serial founder effect is likewise detected using a SNV database independently characterizing worldwide genetic variation ^*41*^. Migration distance is therefore a good proxy for genetic diversity variation. Genetic diversity variation in human populations is the systemic perturbation coupling co-domains in a binary protein complex as investigated here.

Two methods are used to estimated migration distances on earth’s surface. The first method uses great circle distances between waypoints that follow pathways described by Ramachandran et al. (2005) ^*22*^ but with some waypoint additions. The second method uses driving distances between the same waypoints where feasible. The former is shorter than the latter without exception. Together they form informal lower and upper bounds to migration distance estimates that were adjusted incrementally outward from the midpoint of the informal lower and upper bounds, and populations reordered, until a line could be fitted through population migration distances within the new limits. Limits were factors of 1.12 and 1.14 larger than their initial estimates for populations listed in **SI Tables S7 & S8**. Populations in **SI Tables S7 & S8** are subsets of the human populations in **SI Tables S2 & S5** by eliminating those for which distance ranking is not feasible (GLO, OTH) or migration distance redundant (TWC & PCC are both from people in the UK and only PCC was included because it is the larger dataset). Distance estimates (and genetic diversity variation) from the fitted line for populations in **SI Tables S7 & S8** are equally space over the Population Index parameter as needed for the co-domain 2D correlation genetics formalism.

**Fig 5** summarizes migration pathways starting from Addis Ababa, Ethiopia and ending at points used to estimate migration distance for populations listed in **SI Tables S7 & S8**. Migration to all destinations are via Cairo, Egypt except those ending in Africa. Europe is reached from Cairo by two pathways, one via Istanbul, Turkey then west (Cairo to Istanbul distance estimated by driving distance) and, second from Asia after a northerly detour east of the Caspian Sea via Chelyabinsk, Russia. East Asia is reached by 3 routes, two north and one south of the Himalayan mountains. North and South America are reached via a fourth Asian pathway through northern Russia and by more recent forced or voluntary immigration from Africa, Europe, and China. Oceania is reached by the south Asian pathway and by immigration from England. Forced or voluntary immigration are not indicated in the map (see next paragraph).

**Figure 5.**
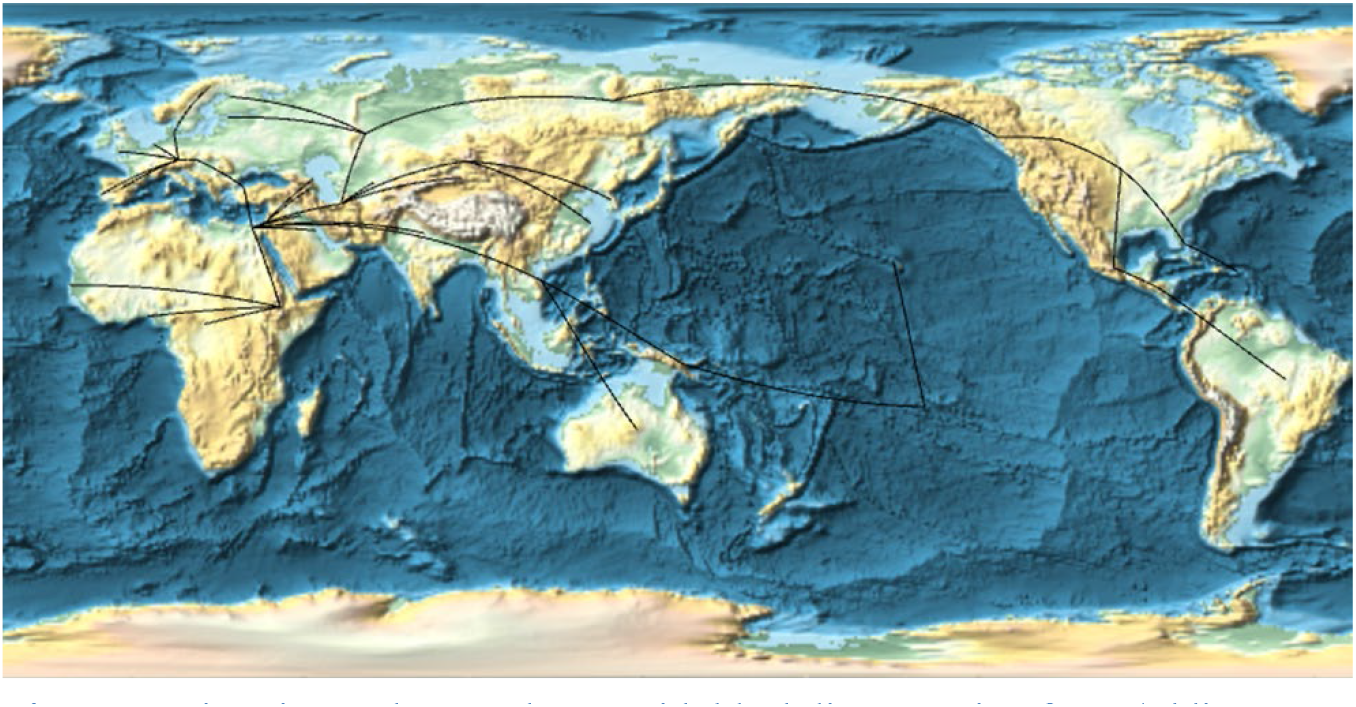
Migration pathways shown with black lines starting from Addis Ababa, Ethiopia and ending at points used to estimate migration distance for populations listed in **SI Tables S7 & S8**. Migration to all destinations, excluding those ending in Africa, are via Cairo, Egypt.

Individuals sampled in North America, South America, and Australia contributed SNV data to the NCBI database that are from native and non-native populations. Overall migration distance to these regions was estimated using a weighted combination of native and non-native path distance with weights proportional to overall ethnic contributions to local population. Two or more race (mixed) individuals contribute fractional parts to each represented single race population (Black Africans, Caucasian, Asian, or Indigenous). Non-native path distance includes just the distance from Addis Ababa to their place of origin before immigration. Native path distance is from Addis Ababa to the destination. Native Hawaiians have a waypoint in the Pacific Ocean at Tahiti. Migration destination for calculating distance for enslaved Black Africans taken to North or South America was Senegal in West Africa.

## 3. RESULTS

Neural/Bayes network models for transduction (**Fig 3**) applied to complex βmys/MYBPC3 or control complex MAPT/MYBPC3 investigates intra-protein contacts. Probability for a SNV from a given population (populations listed in **SI Tables S2 or S5**) falling into domains represented in **Figs 1& 2** (domains listed in **SI Tables S1 or S4**) are computed using eqs. 3-4 for pathogenic or benign outcomes and 20 independent best-of-the-best implicit models for energy transduction (βmys/MYBPC3) or unknown function (MAPT/MYBPC3) in the complex. Transduction models selected satisfy the reproducibility and significance testing described in METHODS (*2*.*3 Neural network validation*). Synchronous and asynchronous 2D correlations (eqs. 5-7) for co-domains are identified by their sensitivity to human population using 2D correlation genetics.

### 3.1 Complex βmys/MYBPC3

**Fig 6** shows 2D correlation genetics maps for complex βmys/MYBPC3 with pathogenic outcomes computed using eqs. 5-7. Axes represent βmys (M7 1-39) followed by MYBPC3 (C3 40-65) domains where M7 and C3 abbreviates βmys and MYBPC3, respectively. Domain name, two letter code, protein sequence, and index number are from **SI Table S1**. Two-letter codes label several protein domains on the leftmost axis in the figure. The grayscale above each 2D map indicates numerical intensities (z-values). Correlation maps divide into 4 regions labeled M7-M7, M7-C3, C3-C3, and C3-M7 that separate inter-protein cross-peaks (M7-M7 and C3-C3) from intra-protein cross-peaks (M7-C3 and C3-M7). Synchronous (left) or asynchronous (right) maps show population coincidental or population sequential changes in SNV probability for domains in the complex. Intensity peaks along the diagonal in the synchronous map are autocorrelated probabilities for each domain. The 6 most significant pathogenic co-domain interactions have combined synchronous and asynchronous co-domain interaction cross-correlates (eq. 8) that are >6.3 standard deviations from the mean. These co-domains are linked by correlation squares (white, green, red, blue, representing different βmys domains then repeating color sequence as needed) that link the off-diagonal interacting domain coordinates falling within M7-C3 and C3-M7 regions. M7-C3 and C3-M7 regions are related by inversion through the diagonal line.

**Figure 6.**
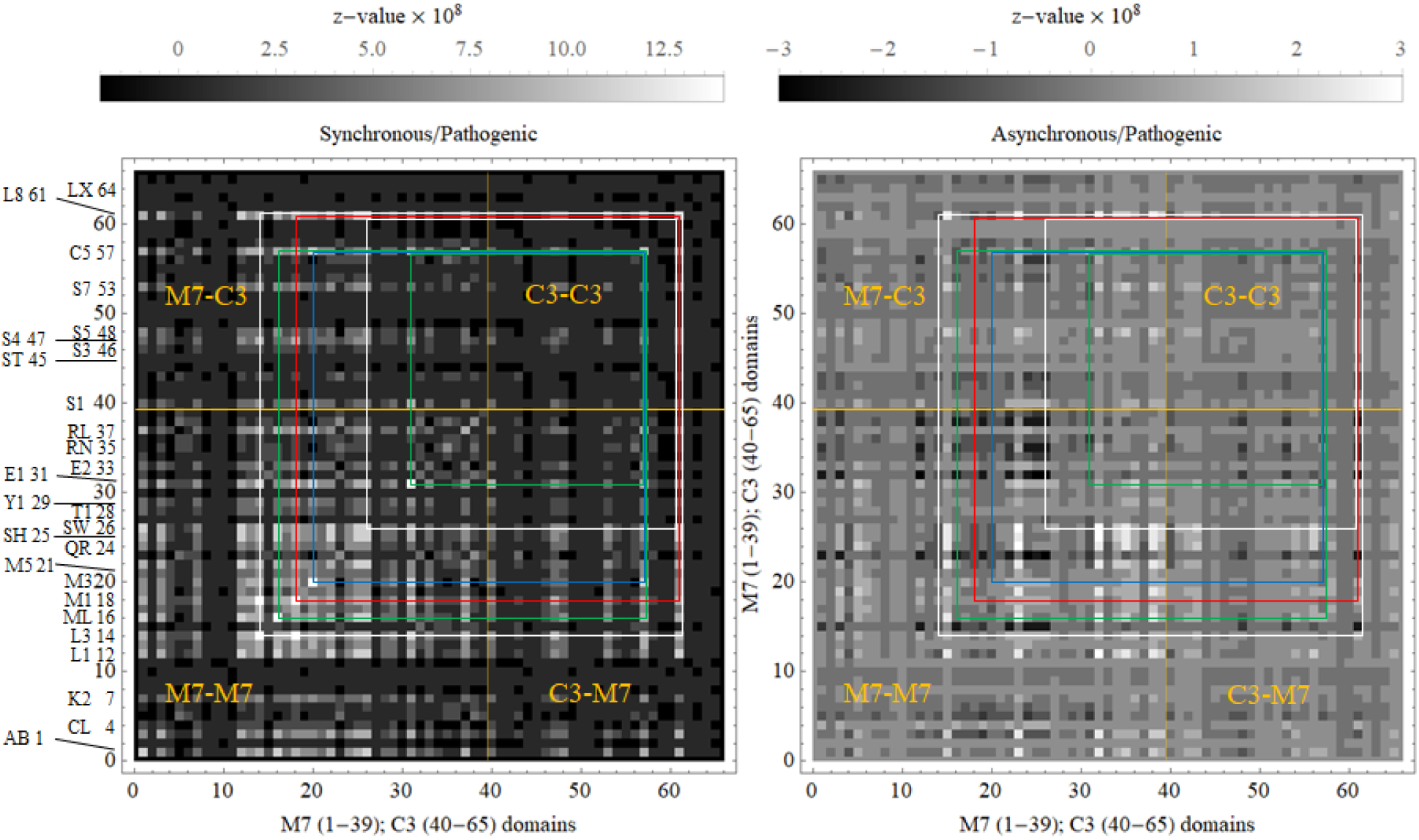
2D correlation genetics maps for complex βmys/MYBPC3 with pathogenic outcomes. Axes represent βmys (M7 1-39) followed by MYBPC3 (C3 40-65) domains. Domain index on the axes is linked to its two letter code and protein sequence in **SI Table S1**. Several protein domains are also labeled by their two-letter code on the leftmost axis. Intensities (z-values) are indicated numerically by the grayscale. The 4 regions defined by vertical and horizontal orange lines at the interface of pixels 39-40 labeled M7-M7, M7-C3, C3-C3, and C3-M7 separate inter-protein cross-peaks (M7-M7 and C3-C3) from intra-protein cross-peaks (M7-C3 and C3-M7). The 6 most significant pathogenic co-domains are linked by correlation squares (white, green, red, blue, for different βmys domains then repeating color sequence as needed) that link the off-diagonal interacting domain coordinates falling within M7-C3 and C3-M7 regions.

**Fig 7 panel a** shows the M7-C3 region for synchronous and asynchronous maps and the portion of the correlation squares falling within them taken from **Fig 6. Fig 7 panel b** red lines represent the co-domain interactions linking βmys and MYBPC3 for both synchronous and asynchronous correlates with the arrow indicating asynchronous pathway phase where leading or lagging (at the pointy end) is relative to the population sequence from **SI Table S7**. Synchronous pathways are also directional but represent simultaneously increasing SNV probabilities in the co-domains for all cases in this study. The paths identify intra-protein transduction pathways interrupted by pathogenic SNV’s. Highest significance synchronous and asynchronous pathways involve MYBPC3 linker 8 (L8) or Ig-like domain 5 (C5) and six diverse βmys domains. L8 leads asynchronous interactions with βmys switch 2 helix (SW), actin binding loop 3 (L3), and the βmys-S2 binding site for MYBPC3-C1 (M1). C5 leads asynchronous interaction with βmys myopathy loop (ML), and, lags asynchronous interactions with ELC EF1 domain (E1) and the βmys-S2 binding site for MYBPC3-C2 (M3). SNVs contributing to the 2D map are pathogenic and affect either the M7 or the C3 side at the ends of the red lines. It suggests that co-domains coordinate function during contraction using the real-or virtual-mechanisms illustrated in **Fig 4**.

**Figure 7 Panel a.**
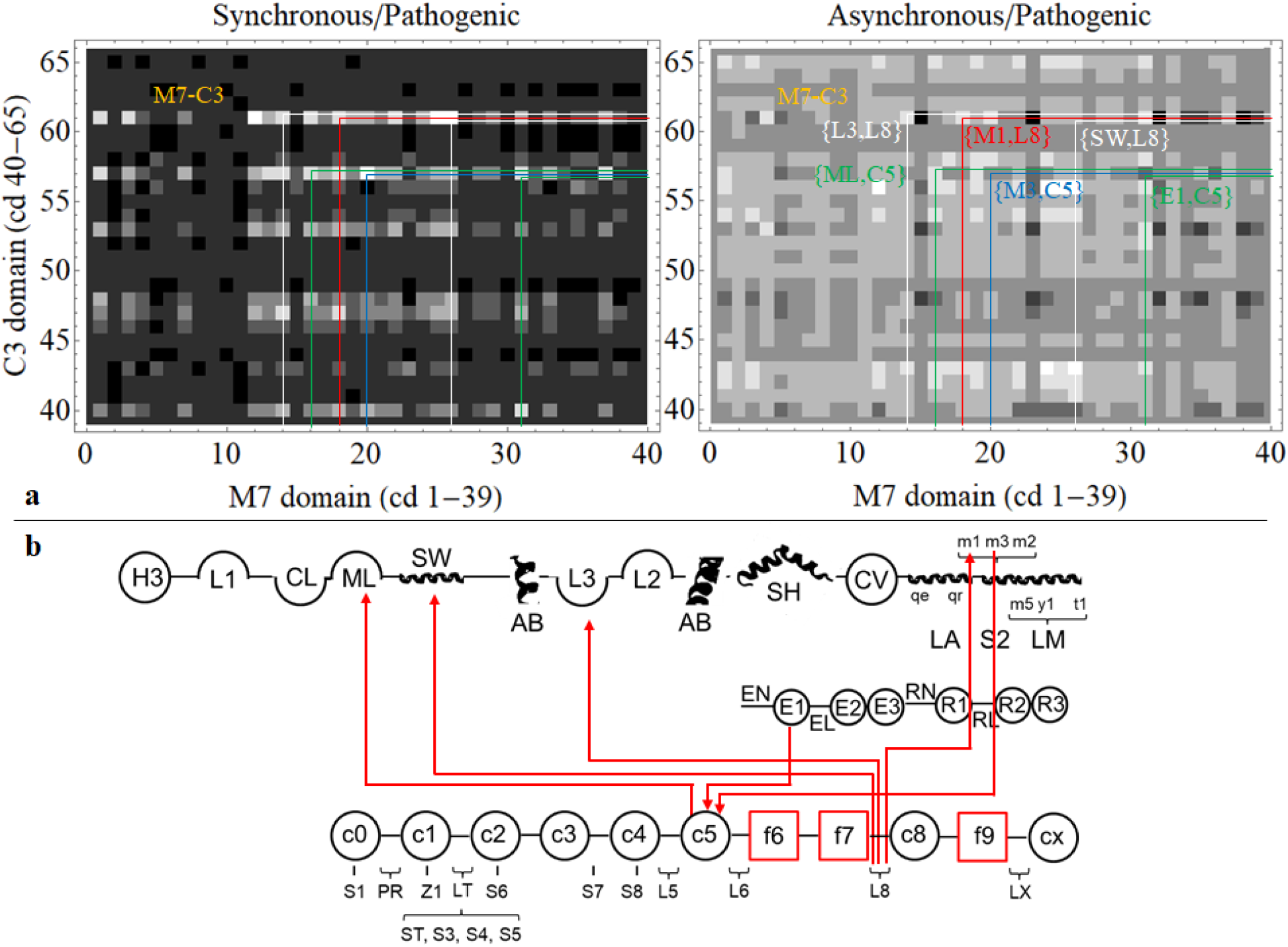
M7-C3 region for synchronous and asynchronous maps and the portion of the correlation squares falling within them for pathogenic outcomes (**Fig 6**). **Panel b**. Red lines show directed co-domain interactions between βmys and MYBPC3. They represent both synchronous and asynchronous correlates with the arrow in each path indicating asynchronous pathway phase where leading or lagging (at the pointy end) is relative to the population sequence from **SI Table S7**.

The L8 domain in MYBPC3 has co-dependence with multiple sites on βmys suggesting the virtual-mechanism wherein a SNV altering the co-domain in MYBPC3 perturbs multiple intra-protein contacts between MYBPC3 and βmys. Hypothetically L8 functions as a hinge in MYBPC3. The same logic applies to C5 where its co-dependence with multiple sites on βmys likewise suggests the virtual-mechanism. The C5 domain in MYBPC3 was already identified as a hinge ^*17*^ supporting the virtual co-domain hypothesis.

**Fig 8 panel a** shows the M7-C3 region for synchronous and asynchronous maps and the portion of the correlation squares falling within them paralleling **Fig 7** but for benign outcomes. **SI Fig S1** shows the full 2D correlation maps for benign outcomes. The 6 most significant co-domain interactions have combined synchronous and asynchronous co-domain interaction cross-correlates (eq. 8) that are >3.7 standard deviations from the mean. **Fig 8 panel b** red lines show the co-domain interactions linking βmys and MYBPC3 representing both synchronous and asynchronous correlates with the arrow indicating asynchronous pathway phase where leading or lagging (at the pointy end) is relative to the population sequence from **SI Table S7**. Highest significance synchronous and asynchronous pathways involve MYBPC3 phosphorylatable serines S1(Ser47), ST(Ser273), and S3 (Ser282) and three domains of βmys. ST and S3 in the MYBPC3 regulatory domain and S1 lead asynchronous interactions with βmys actin binding C-loop (CL). ST and S3 lead asynchronous interactions with βmys-LM titin binding site (T1 ^*29, 31*^). S3 lags an asynchronous interaction with βmys-LM MYBPC3-CX binding site (M5 ^*29*^). SNVs contributing to the 2D map are benign and affect either the M7 or the C3 side at the ends of the red lines. It suggests that co-domains coordinate function during contraction using the real-or virtual-mechanisms illustrated in **Fig 4**.

**Figure 8 Panel a.**
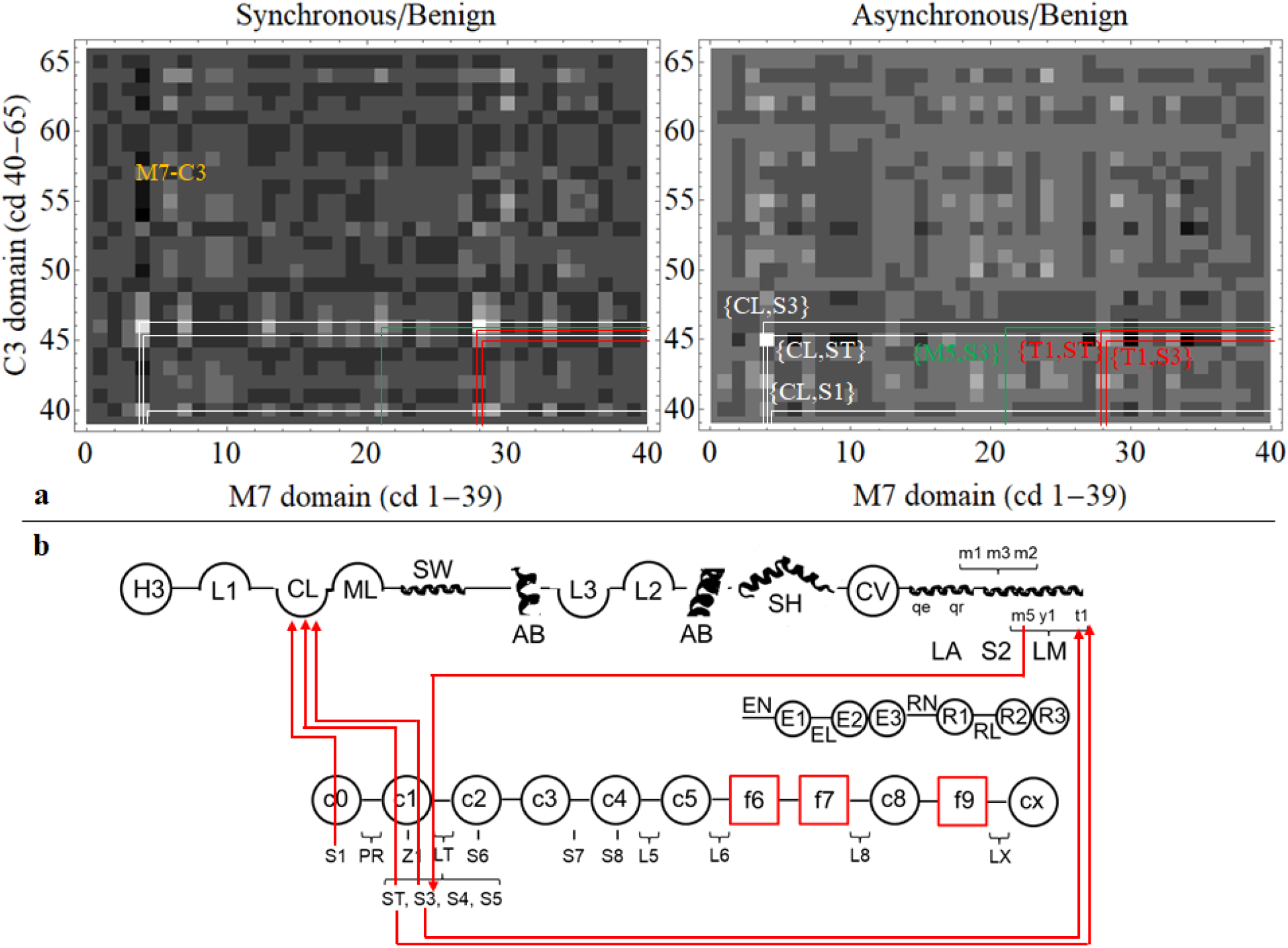
M7-C3 regions for synchronous and asynchronous maps and the portion of the correlation squares falling within them for benign outcomes all taken from **SI Fig S1**. Nomenclature is otherwise identical to **Fig 7 panel a. Panel b**. Red lines show directed co-domain interactions between βmys and MYBPC3. Nomenclature is otherwise identical to **Fig 7 panel b**.

The actin binding C-loop domain in βmys regulates actin-activated myosin ATPase ^*42*^ and modulates contraction velocity in coordination with actin bound tropomyosin ^*43*^. It engages the MYBPC3 regulatory domain in linker 2 (LT) and S1 in C0 suggesting joint βmys/MYBPC3 regulation of actin-activated myosin ATPase and actomyosin translation speed. Other co-domains identified involve MYBPC3 sites in LT interacting with βmys sites in LM mostly near the βmys C-terminus recalling the transient binding of the MYBPC3 N-terminal peptide C0-C1-LT with the thick filament ^*44*^.

The evidence suggests pathogenic vs benign co-domains selectively identify mechanical (**Fig 7b**) vs regulatory (**Fig 8b**) transduction functions within the complex. It appears co-domains in βmys linked to its mechanical function are less resilient to SNV modification and become less reliable with decreasing genetic diversity, i.e., migration out of Africa.

Cross-peak intensity averaged over the 6 most significant co-domains identified by pathogenic or benign SNV’s illustrate co-domain population dependence. Averaged quantities are defined in eqs. 9-10 for the {α,ε} ≡{βmys,MYBPC3} co-domains where pathogenic {α,ε} = {ML,C5}, {M3,C5}, {E1,C5}, {SW,L8}, {L3,L8}, and {M1,L8} or benign {α,ε} = {CL,S1}, {CL,ST}, {CL,S3}, {M5,S3}, {T1,ST}, and {T1,S3}. **SI Fig S2 panel a** indicates synchronous interactions for pathogenic (black) or benign (blue) SNV’s and **panel b** the asynchronous interactions for pathogenic (left) or benign (right) SNV’s. Population three letter code and index are indicated in **panel a**. Asynchronous population dependence (**panel b**) involves population pairs requiring two dimensions for display.

Highest amplitude synchronous correlations (**panel a**) for pathogenic SNV’s include African (AFR), Ashkenazi Jewish (ASJ), European (EUR), Asian (ASI), and American (AMR) populations. Lowest amplitude synchronous correlations for pathogenic SNV’s include European American (EUA) and African American (AFM) populations. Highest and lowest amplitude synchronous correlations for benign SNV’s follow a similar pattern as pathogenic SNV’s but with less dynamic range. The high amplitude synchronous correlations fall into two groups at the highest (3,800-5,500 km) and middle range (9,000-10,000 km) genetic differentiation categories.

Highest amplitude asynchronous correlations for pathogenic SNV’s (**panel b left**) couple AFR⟶ASJ, ASJ⟶EUR, and AMR⟶ASI. They couple the highest amplitude synchronous correlations and indicate direction with oppositely signed peaks relative to the diagonal. Highest amplitude asynchronous correlation for benign SNV’s (**panel b right**) couple AFR⟶ASJ.

### 3.2 Control complex MAPT/MYBPC3

The analysis just described for βmys/MYBPC3 was also performed on control complex MAPT/MYBPC3. **SI Figs S3 & S4** show 2D correlation results for MAPT/MYBPC3 for pathogenic and benign outcomes and paralleling the 2D correlation results for βmys/MYBPC3 in **Fig 6** and **SI Fig S1. Figs 9 & 10 panels a** show the C3-MT regions for the synchronous and asynchronous maps, the portion of the correlation squares falling within them, and the list of domain pairs corresponding to the 6 most significant co-domain cross-peaks for pathogenic and benign outcomes. These data are summarized by the co-domain interactions in **Figs 9 & 10 panels b** indicating potential intra-protein transduction pathways infiltrated by pathogenic or benign SNVs. Standard deviations >4.7 and >2.3 from their mean admit the 6 most significant pathogenic and benign co-domain cross-correlates. Pathogenic and benign thresholds for 0 co-domains in complex MAPT/MYBPC2 are 5.4 and 2.7 standard deviations from the mean compared to 6.6 and 6.1 in complex βmys/MYBPC3 implying no false positives in the βmys/MYBPC3 complex. Standard deviations of 1.2 and 3.4 separate the system with known intra-protein contacts from the one with (presumably) none suggesting the statistical headroom for distinguishing them. The benign species shows the larger contrast.

**Figure 9 Panel a.**
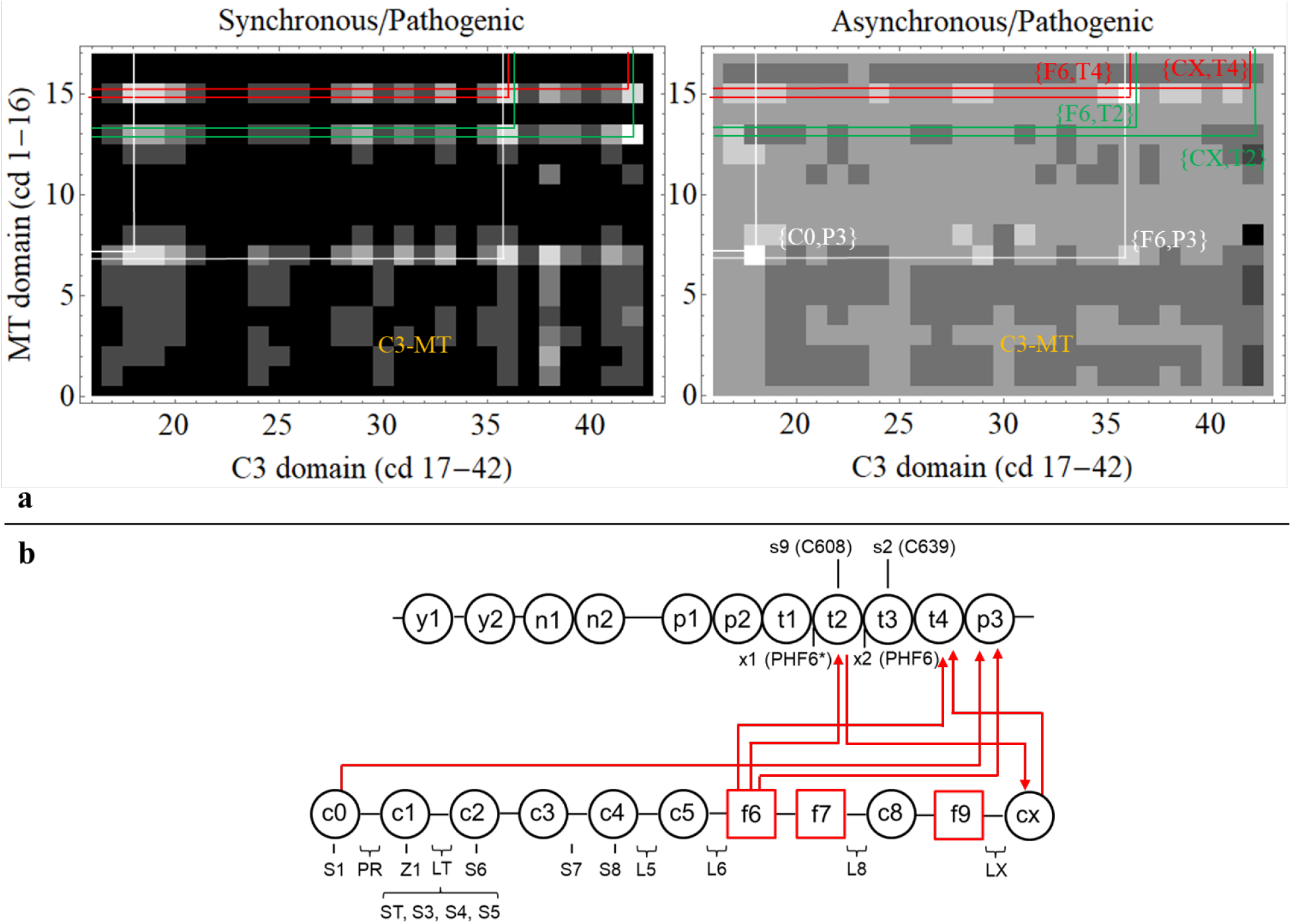
C3-MT region for synchronous and asynchronous maps and the portion of the correlation squares falling within them for pathogenic outcomes all taken from **SI Fig S3. Panel b**. Red lines show directed co-domain interactions between MAPT and MYBPC3. They represent both synchronous and asynchronous correlates with the arrow in each path indicating asynchronous pathway phase where leading or lagging (at the pointy end) is relative to the population sequence from **SI Table S8**. Synchronous pathways are also directional but represent simultaneously increasing SNV probabilities in the co-domains for all cases.

## Discussion

Cardiac muscle sarcomere proteins comprise the molecular machinery generating and regulating muscle contraction. Their coordinated action involves most critically the βmys motor powering contraction by ATP free energy transduction into mechanical work and MYBPC3 in a phosphorylation dependent regulatory role. This contractile system subset dynamically adapts contractile force and velocity using myosin step-size modulation ^*45, 46*^ and a coordinated βmys/MYBPC3 interaction ^*11*^. Their significance in cardiac muscle contraction is affirmed by the observation that familial hypertrophic cardiomyopathy is most often associated with mutation in one of these two proteins ^*47*^.

Structural information pertaining to adaptive contractile force and velocity is often static and at highest resolution when individual sarcomere proteins are crystallized ^*48*^ or at usually lower resolution when larger reconstituted systems are frozen for cryoEM ^*49*^. 2D-NMR structures include dynamic characteristics by using structural ensembles generated by constrained molecular dynamics simulation ^*50*^. In vivo approaches include single zebrafish skeletal myosin motor dynamics in isometric contraction ^*51*^ and βmys motor step-size dynamics in the beating zebrafish heart ^*45*^. Introduced here is a computational approach involving worldwide human genetics data from hearts in vivo. Nonsynonymous SNV residue substitutions are the probes that modify βmys or MYBPC3 in the human heart. The NCBI SNP database containing results from this large-scale natural experiment consistently records SNV physical characteristics including substituted residue location in the protein functional domain, side chain substitution, substitution frequency, and human population group, but inconsistently records SNV implicated phenotype and pathology outcomes. A consistent subset of the data trains and validates a feed-forward neural network modeling the contraction mechanism. The full database is completed using the model then interpreted probabilistically with a discrete Bayes network to provide the SNV probability for a functional domain location given pathogenicity and human population ^*19, 21*^. Applying the approach to complex βmys/MYBPC3 identifies potential intra-protein pathways coupling these key proteins in the sarcomere pertaining to muscle shortening power generation and regulation in the human heart. The intra-protein pathways involve functional domains coupling the two proteins called co-domains. They are identified by their cross-correlated SNV probability responses to the perturbation provided by the divergent human genetics in different populations using 2D correlation genetics. It is analogous to the 2D NMR study of protein complex structure indicated by the cross-correlated response of specific nuclei to the perturbation provided by judiciously selected radio frequency (RF) pulse sequences and magnetization transfer. Co-domain identification in a multiprotein complex as described here implies a potential to estimate spatial proximity constraints in dynamic protein interactions in vivo via structural ensembles elaborated by constrained molecular dynamics simulation also analogous to 2D-NMR based structure determination.

In this study, βmys/MYBPC3 and MAPT/MYBPC3 interactions were investigated. The former is known to complex and participate in force/velocity generation and regulation in a working heart. The latter is unlikely to meet in vivo hence its relevance as a control. Cross-protein functional domains correlated by human population in 2D identify co-domains in the βmys/MYBPC3 complex as shown in **Figs 6 & 7** for pathogenic SNV’s. Pathogenic SNV’s alter gene product protein structure to impair functional pathways through co-domains. The pathogenic case implies co-domains cannot maintain functional integrity when modified over the shifting genetic context of the human population index. The 6 highest significance co-domains for the complex are identified as the cross-peaks in correlation squares for synchronous and asynchronous interactions. Synchronous/asynchronous nomenclature refers here to genetic diversity ordered human populations (**SI Tables S7 & S8**). It identifies domain pairs whose SNV probabilities synchronize with, lead, or lag each other over populations ordered by quantitation of their genetic diversity. Leading domain SNV probabilities are more affected by populations containing higher genetic diversity vs the lagging domain where SNV probabilities are more impacted by populations containing lower genetic diversity.

The focus on co-domain coupling is emphasized by **Figs 7 panel a** where cross-peaks linking βmys with MYBPC3 (the M7-C3 region in **Fig 6**) are shown. These data are interpreted by co-domain maps in **Fig 7 panel b** for combined synchronous and asynchronous interactions of pathogenic SNVs. It shows 3 directed pathways linking L8 in MYBPC3 with actin binding (L3), energy transduction (SW), and C1 binding (M1) ^*15*^ related functional domains in βmys. The peptide linker in the MYBPC3 N-terminus between C1 and C2 (linker 2 or LT) has phosphorylation sites participating in contractile regulation by modulating myosin activity ^*9-11*^ and calcium sensitivity ^*12*^. It is reasonable to assume that disrupting C1 binding to βmys could impact the regulatory function on contraction exerted by LT. The L8-βmys interaction is the model for the virtual co-domain mechanism in **Fig 4** wherein a SNV altering structure in MYBPC3 L8 affects coupling to multiple real co-domains in βmys.

**Fig 7 panel b** also shows 3 directed pathways linking SNVs in C5 from MYBPC3 with actin binding (ML), force generation (E1), and C2 binding (M3) ^*15*^ related functional domains in βmys. Like with L8, it is interpreted as indicating SNVs in C5 disrupt multiple co-domain interactions involving ML, E1, and the regulatory domain (LT) between C1 and C2. It again implies the virtual co-domain effect but involving C5 wherein a SNV altering structure in MYBPC3 C5 affects coupling to multiple real co-domains in βmys.

MYBPC3 LT and C5 hinges were identified ^*17*^ independently corroborating the hypothesis that SNVs in C5 can disrupt bending in the MYBPC3 that is key to forming real co-domain interactions in complex βmys/MYBPC3. A similar role proposed for linker 8 (L8) implies it is a third flexible linker. It is near the C-terminus and would facilitate the multiple co-domain interactions proposed for the βmys/MYBPC3 complex while the MYBPC3 C-terminus is bound to LM on the thick filament.

**Fig 8** shows the subset of cross-peaks linking βmys with MYBPC3. It is equivalent to **Fig 7** except for benign SNVs. The 6 highest significance co-domains for the complex are identified as the cross-peaks in correlation squares for synchronous and asynchronous interactions. Benign SNV’s alter product sequence without impairing function of resilient domains correlated by human population. The benign case implies co-domains maintain functional integrity when modified even over the shifting genetic context of the population index (**SI Table S7**). **Fig 8 panel a** shows cross-peaks linking βmys with MYBPC3 (the M7-C3 region in **SI Fig S1**). These data are interpreted as the co-domain map in **Fig 8 panel b** for combined synchronous and asynchronous interactions of benign SNV’s. It shows directed pathways link SNVs in βmys at the C-loop (CL) actin binding site to phosphorylation sites in the regulatory domain of MYBPC3 at LT. It implies and confirms the regulatory control mechanism at LT in MYBPC3. It also implies participation of the C-loop in βmys/MYBPC3 modulated force/velocity regulation ^*52*^. The C-loop is known to participate in energy transduction ^*42*^, actin-activation of myosin ATPase ^*53*^ and to modulate myosin in vitro motility velocity in the presence of actin bound tropomyosin ^*43*^. The latter interaction was confirmed in a static structure of a thick filament ^*54*^. It seems that the C-loop maintains resilient links with the MYBPC3 regulatory domain.

**Fig 8 panel b** also shows directed pathways involve SNVs in βmys at LM and in MYBPC3 at the regulatory domain LT. LM sites at M5 and T1 engage with ST, S3, or both. M5 and T1 are binding sites for CX (MYBPC3) and titin ^*29, 31*^, respectively. It implies that several sites on LM maintain interactions with S3, a principal phosphorylatable serine in the MYBPC3 regulatory domain. The βmys C-loop and LM maintain robust lines of communication with the MYBPC3 regulatory domain that stand despite SNV modification and under the changing genetic background of human populations.

The evidence suggests pathogenic vs benign co-domains selectively identify mechanical vs regulatory transduction functions within the complex. It appears co-domains in βmys linked to its mechanical function are less resilient to SNV modification and become less reliable with decreasing genetic diversity, i.e., migration out of Africa.

The analysis described for βmys/MYBPC3 is applied to the control complex MAPT/MYBPC3. These data are summarized in **SI Figs S3 & S4** and **Figs 9 & 10**. Pathogenic and benign thresholds for 0 co-domains in complex MAPT/MYBPC2 are 5.4 and 2.7 standard deviations compared to 6.6 and 6.1 in complex βmys/MYBPC3. The contrasting thresholds imply there are no false positives for co-domains in complex βmys/MYBPC3. Minima of 1.2 and 3.4 standard deviations separate systems with known intra-protein contacts from one with (presumably) none suggesting the statistical headroom for distinguishing them.

**Figure 10 Panel a.**
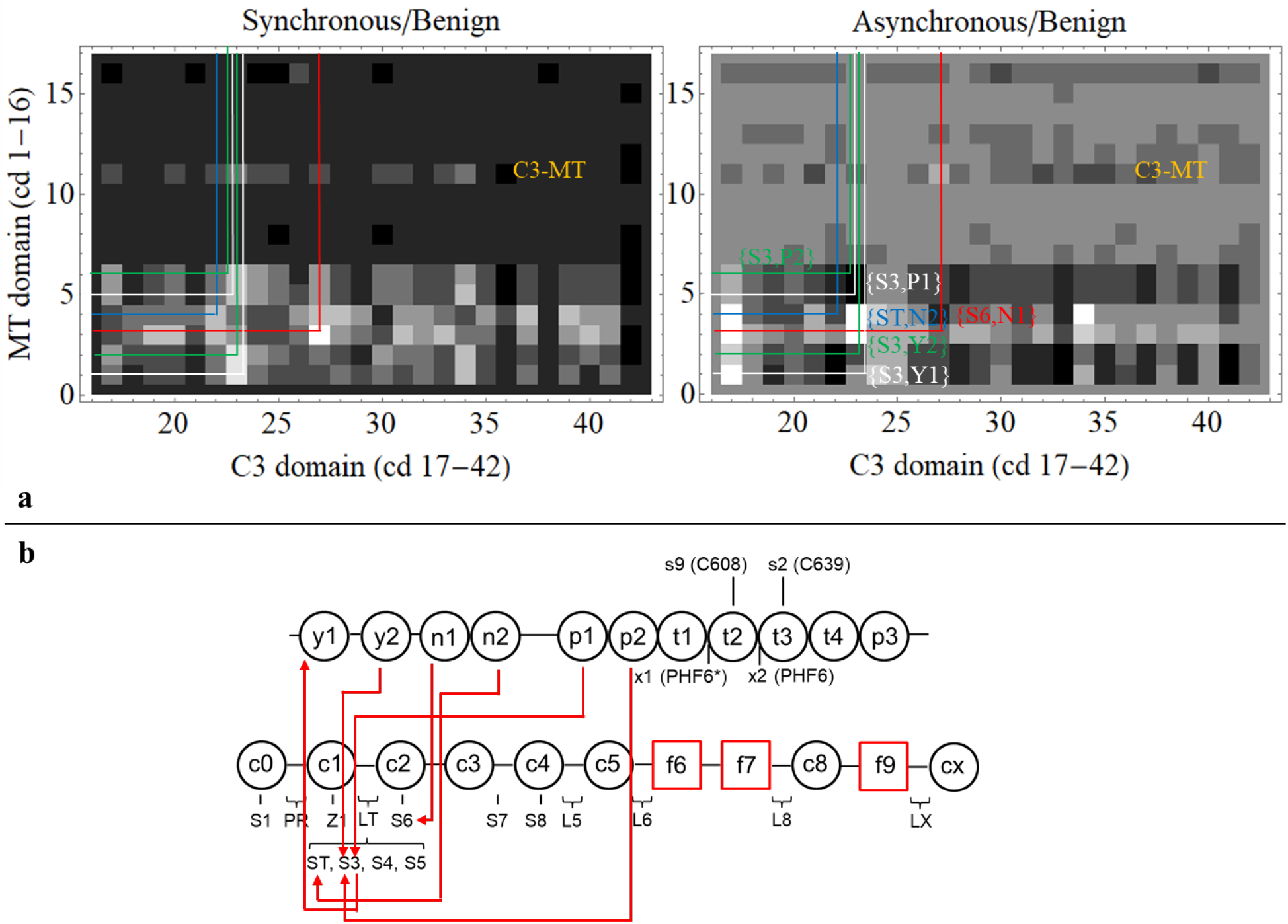
MT-C3 regions for synchronous and asynchronous maps and the portion of the correlation squares falling within them for benign outcomes all taken from **SI Fig S4**. Nomenclature is otherwise identical to **Fig 8 panel a. Panel b**. Red lines show directed co-domain interactions between MAPT and MYBPC3. Nomenclature is otherwise identical to **Fig 8 panel b**.

## Conclusion

βmys and MYBPC3 in complex is a minimal model for the contractile system in the heart wherein dynamic intra-protein interactions for system regulation are ascertained. The database from a worldwide in vivo experiment collecting SNVs in the two proteins consistently records their physical characteristics including substituted residue location in the protein functional domain, side chain replacement, substitution frequency, and human population group, but inconsistently records SNV implicated phenotype and pathology outcomes. A consistent subset of the data trains and validates a feed-forward neural network modeling the contraction mechanism. The full database is completed using the model then interpreted probabilistically using a discrete Bayes network providing the SNV probability for pathogenicity given the functional domain location and human population. Co-domains, intra-protein domains coupling βmys and MYBPC3, are identified by their population correlated SNV probability product for given pathogenicity. Divergent genetics in the human populations couple co-domains in this method called 2D correlation genetics. Pathogenic and benign SNV data implies three co-domain hubs, C5 and L8, in MYBPC3 with links to several domains across the βmys motor, and, C-loop (CL) in βmys with links to the MYBPC3 regulatory domain (LT). C5 (MYBPC3) links with actin binding, force generation, and C0-C2 binding sites in βmys. L8 (MYBPC3) links with actin binding, energy transduction, and C0-C2 binding sites in βmys. These critical sites in MYBPC3 are known (C5) and proposed (L8) regions that bend. The critical site in βmys (CL) is an actin binding site, an actin contact sensor regulating actin-activated myosin ATPase, and a contraction velocity modulator related to its interaction with actin bound tropomyosin. Links between CL (βmys) and LT (MYBPC3) impact the principal functions of the cardiac contractile system. The identification of co-domains in a multiprotein complex implies a potential to estimate spatial proximity constraints for dynamic protein interactions in vivo. These can be modeled via structural ensembles elaborated by constrained molecular dynamics simulation in a process analogous to 2D-NMR protein structure determination.

## Supporting information

6ddpbmysMYBPC3

6ddpMAPTMYBPC3

## Acknowledgment

The author thanks Katalin Ajtai for scientific discussion and critical review of the manuscript.

## Supplementary Information

Supplementary information (SI) consists of tables: **Tables S1-S8**, and figures: **Figures S1-S4**, and data sets for the fulfilled and unknown 6ddps for complex βmys/MYPBC3 and MAPT/MYBPC3. Data sets are contained in files 6ddpbmysMYBPC3.xls and 6ddpMAPTMYBPC3.xls.

## References

[1] Al-Khayat, H. A., Kensler, R. W., Squire, J. M., Marston, S. B., and Morris, E. P. (2013) Atomic model of the human cardiac muscle myosin filament, Proc. Natl. Acad. Sci. USA 110, 318–323.

[2] Lorenz, M., and Holmes, K. C. (2010) The actin-myosin interface, Proc. Nat. Acad. Sci. USA 107, 12529–12534.

[3] Highsmith, S., and Eden, D. (1993) Myosin-ATP chemomechanics, Biochemistry 32, 2455–2458.

[4] Rayment, I., and Holden, H. M. (1993) Myosin subfragment-1: structure and function of a molecular motor, Curr. Opin. Struct. Biol. 1993, 944–952.

[5] Lowey, S., Waller, G. S., and Trybus, K. M. (1993) Function of skeletal muscle myosin heavy and light chain isoforms by an in vitro motility assay, J. Biol. Chem. 268, 20414–20418.

[6] Greenberg, M. J., Kazimierczak, K., Szczesna-Cordary, D., and Moore, J. R. (2010) Cardiomyopathy-linked myosin regulatory light chain mutations disrupt myosin strain-dependent biochemistry, Proc. Natl. Acad. Sci. USA 107, 17403–17408.

[7] Yadav, S., Kazmierczak, K., Liang, J., Sitbon, Y. H., and Szczesna-Cordary, D. (2018) Phosphomimetic-mediated in vitro rescue of hypertrophic cardiomyopathy linked to R58Q mutation in myosin regulatory light chain, The FEBS Journal 286, 151–168.

[8] Previs, M. J., Previs, S. B., Gulick, J., Robbins, J., and Warshaw, D. M. (2012) Molecular Mechanics of Cardiac Myosin-Binding Protein C in Native Thick Filaments, Science 337, 1215–1218.

[9] Kensler, R. W., Craig, R., and Moss, R. L. (2017) Phosphorylation of cardiac myosin binding protein C releases myosin heads from the surface of cardiac thick filaments, Proc. Natl. Acad. Sci. USA 114, E1355–E1364.

[10] McNamara, J. W., Singh, R. R., and Sadayappan, S. (2019) Cardiac myosin binding protein-C phosphorylation regulates the super-relaxed state of myosin, Proc. Natl. Acad. Sci. USA 116, 11731–11736.

[11] Nelson, S. R., Li, A., Beck-Previs, S., Kennedy, G. G., and Warshaw, D. M. (2020) Imaging ATP Consumption in Resting Skeletal Muscle: One Molecule at a Time, Biophys. J. 119, 1050–1055.

[12] Inchingolo, A. V., Previs, S. B., Previs, M. J., Warshaw, D. M., and Kad, N. M. (2019) Revealing the mechanism of how cardiac myosin-binding protein C N-terminal fragments sensitize thin filaments for myosin binding, Proc. Natl. Acad. Sci. USA 116, 6828–6835.

[13] Craig, R., Lee, K. H., Mun, J. Y., Torre, I., and Luther, P. K. (2014) Structure, sarcomeric organization, and thin filament binding of cardiac myosin-binding protein-C Pflugers Arch - Eur J Physiol 466, 425–431.

[14] Rahmanseresht, S., Lee, K. H., O’Leary, T. S., McNamara, J. W., Sadayappan, S., Robbins, J., Warshaw, D. M., Craig, R., and Previs, M. J. (2021) The N terminus of myosin-binding protein C extends toward actin filaments in intact cardiac muscle, J. Gen. Physiol. 153.

[15] Ratti, J., Rostkova, E., Gautel, M., and Pfuhl, M. (2011) Structure and Interactions of Myosin-binding Protein C Domain C0: Cardiac-Specific Regulation of Myosin at Its Neck?, In J. Biol. Chem., pp 12650–12658.

[16] Luther, P. K., Winkler, H., Taylor, K., Zoghbi, M. E., Craig, R., Pedron, R., Squire, J. M., and Liu, J. (2008) Direct visualization of myosin-binding protein C bridging myosin and actin filament in intact muscle, Proc. Nat. Acad. Sci. USA 108, 11423–11428.

[17] Previs, M. J., Mun, J. Y., Michalek, A. J., Previs, S. B., Gulick, J., Robbins, J., Warshaw, D. M., and Craig, R. (2016) Phosphorylation and calcium antagonistically tune myosin-binding protein C’s structure and function, Proc. Natl. Acad. Sci. USA 113, 3239–3244.

[18] Harris, S. P. (2021) Making waves: A proposed new role for myosin-binding protein C in regulating oscillatory contractions in vertebrate striated muscle, J. Gen. Physiol. 153, 1–15.

[19] Burghardt, T. P. (2019) Demographic model for inheritable cardiac disease, Arch. Biochem. Biophys. 672, 108056.

[20] Kraft, T., and Montag, J. (2019) Altered force generation and cell-to-cell contractile imbalance in hypertrophic cardiomyopathy, Pflugers Arch - Eur J Physiol 471, 719–733.

[21] Burghardt, T. P., and Ajtai, K. (2018) Neural/Bayes network predictor for inheritable cardiac disease pathogenicity and phenotype, J Molec Cell Cardiol 119, 19–27.

[22] Ramachandran, S., Deshpande, O., Roseman, C. C., Rosenberg, N. A., Feldman, M. W., and Cavalli-Sforza, L. L. (2005) Support from the relationship of genetic and geographic distance in human populations for a serial founder effect originating in Africa, Proc. Natl. Acad. Sci. U. S. A. 102, 15942–15947.

[23] Noda, I., Dowrey, A. E., Marcott, C., Story, G. M., and Ozaki, Y. (2000) Generalized Two-Dimensional Correlation Spectroscopy, Applied Spectroscopy 54, 236A–248A.

[24] Botts, J., Takashi, R., Torgerson, P., Hozumi, T., Muhlrad, A., Mornet, D., and Morales, M. F. (1984) On the mechanism of energy transduction in myosin subfragment 1, Proc. Natl. Acad. Sci. USA 81, 2060–2064.

[25] Trivedi, D. V., Adhikari, A. S., Sarkar, S. S., Ruppel, K. M., and Spudich, J. A. (2018) Hypertrophic cardiomyopathy and the myosin mesa: viewing an old disease in a new light, Biophysical reviews 10, 27–48.

[26] Woodhead, J. L., and Craig, R. (2020) The mesa trail and the interacting heads motif of myosin II, Arch. Biochem. Biophys. 680, 108228.

[27] Ababou, A., Gautel, M., and Pfuhl, M. (2007) Dissecting the N-terminal Myosin Binding Site of Human Cardiac Myosin-binding Protein C: STRUCTURE AND MYOSIN BINDING OF DOMAIN C2, J. Biol. Chem. 282, 9204–9215.

[28] Ababou, A., Rostkova, E., Mistry, S., Masurier, C. L., Gautel, M., and Pfuhl, M. (2008) Myosin Binding Protein C Positioned to Play a Key Role in Regulation of Muscle Contraction: Structure and Interactions of Domain C1, J. Mol. Biol. 384, 615–630.

[29] Flashman, E., Watkins, H., and Redwood, C. (2007) Localization of the binding site of the C-terminal domain of cardiac myosin-binding protein-C on the myosin rod, The Biochemical journal 401, 97–102.

[30] Obermann, W. M. J., Gautel, M., Weber, K., and Fürst, D. O. (1997) Molecular structure of the sarcomeric M band: mapping of titin and myosin binding domains in myomesin and the identification of a potential regulatory phosphorylation site in myomesin, The EMBO Journal 16, 211–220.

[31] Houmeida, A., Holt, J., Tskhovrebova, L., and Trinick, J. (1995) Studies of the interaction between titin and myosin, J. Cell Biol. 131, 1471–1481.

[32] Consortium, T. U. (2018) UniProt: a worldwide hub of protein knowledge, Nucleic Acids Res. 47, D506–D515.

[33] Weingarten, M. D., Lockwood, A. H., Hwo, S. Y., and Kirschner, M. W. (1975) A protein factor essential for microtubule assembly, Proc. Natl. Acad. Sci. U. S. A. 72, 1858–1862.

[34] Levine, Z. A., Larini, L., LaPointe, N. E., Feinstein, S. C., and Shea, J.-E. (2015) Regulation and aggregation of intrinsically disordered peptides, Proc. Natl. Acad. Sci. USA 112, 2758–2763.

[35] Mazanetz, M. P., and Fischer, P. M. (2007) Untangling tau hyperphosphorylation in drug design for neurodegenerative diseases, Nature Reviews Drug Discovery 6, 464–479.

[36] Nizynski, B., Dzwolak, W., and Nieznanski, K. (2017) Amyloidogenesis of Tau protein, Protein Sci. 26, 2126–2150.

[37] Schweers, O., Mandelkow, E. M., Biernat, J., and Mandelkow, E. (1995) Oxidation of cysteine-322 in the repeat domain of microtubule-associated protein tau controls the in vitro assembly of paired helical filaments, Proc. Natl. Acad. Sci. USA 92, 8463–8467.

[38] von Bergen, M., Friedhoff, P., Biernat, J., Heberle, J., Mandelkow, E.-M., and Mandelkow, E. (2000) Assembly of τ protein into Alzheimer paired helical filaments depends on a local sequence motif (306VQIVYK311) forming β structure, Proc. Natl. Acad. Sci. USA 97, 5129–5134.

[39] von Bergen, M., Barghorn, S., Li, L., Marx, A., Biernat, J., Mandelkow, E.-M., and Mandelkow, E. (2001) Mutations of Tau Protein in Frontotemporal Dementia Promote Aggregation of Paired Helical Filaments by Enhancing Local β-Structure, J. Biol. Chem. 276, 48165–48174.

[40] Deshpande, O., Batzoglou, S., Feldman, M. W., and Cavalli-Sforza, L. L. (2009) A serial founder effect model for human settlement out of Africa, Proc Biol Sci 276, 291–300.

[41] Li, J. Z., Absher, D. M., Tang, H., Southwick, A. M., Casto, A. M., Ramachandran, S., Cann, H. M., Barsh, G. S., Feldman, M., Cavalli-Sforza, L. L., and Myers, R. M. (2008) Worldwide Human Relationships Inferred from Genome-Wide Patterns of Variation, Science 319, 1100–1104.

[42] Ajtai, K., Garamszegi, S. P., Park, S., Velazquez Dones, A. L., and Burghardt, T. P. (2001) Structural characterization of β-cardiac myosin subfragment 1 in solution, Biochemistry 40, 12078–12093.

[43] Ajtai, K., Halstead, M. F., Nyitrai, M., Penheiter, A. R., Zheng, Y., and Burghardt, T. P. (2009) The Myosin C-Loop Is an Allosteric Actin Contact Sensor in Actomyosin, Biochemistry 48, 5263–5275.

[44] Lee, K., Harris, S. P., Sadayappan, S., and Craig, R. (2015) Orientation of Myosin Binding Protein C in the Cardiac Muscle Sarcomere Determined by Domain-Specific Immuno-EM, J. Mol. Biol. 427, 274–286.

[45] Burghardt, T. P., Sun, X., Wang, Y., and Ajtai, K. (2017) Auxotonic to Isometric Contraction Transitioning in a Beating Heart Causes Myosin Step-Size to Down Shift, PLoS ONE 12, e0174690.

[46] Wang, Y., Ajtai, K., and Burghardt Thomas, P. (2018) Cardiac and skeletal actin substrates uniquely tune cardiac myosin strain-dependent mechanics, Open Biology 8, 180143.

[47] Marian, A. J., and Braunwald, E. (2017) Hypertrophic Cardiomyopathy: Genetics, Pathogenesis, Clinical Manifestations, Diagnosis, and Therapy, Circ. Res. 121, 749–770.

[48] Allingham, J. S., Smith, R., and Rayment, I. (2005) The structural basis of blebbistatin inhibition and specificity for myosin II, Nature Structural & Molecular Biology 12, 378–379.

[49] Yamada, Y., Namba, K., and Fujii, T. (2020) Cardiac muscle thin filament structures reveal calcium regulatory mechanism, Nature Communications 11, 153.

[50] Burghardt, T. P., Juranic, N., Macura, S., Muhlrad, A., and Ajtai, K. (1999) Circular Dichroism constrains NMR derived structures of a folded trinitrophenylated hexapeptide in solution, J.Am.Chem.Soc. 121, 10373–10378.

[51] Sun, X., Ekker, S. C., Shelden, E. A., Takubo, N., Wang, Y., and Burghardt, T. P. (2014) In vivo orientation of single myosin lever-arms in zebrafish skeletal muscle, Biophys. J. 107, 1403–1414.

[52] Tanner, B. C. W., Previs, M. J., Wang, Y., Robbins, J., and Palmer, B. M. (2021) Cardiac myosin binding protein-C phosphorylation accelerates β-cardiac myosin detachment rate in mouse myocardium, Am J Physiol Heart Circ Physiol.

[53] Ajtai, K., Garamszegi, S. P., Watanabe, S., Ikebe, M., and Burghardt, T. P. (2004) The myosin cardiac loop participates functionally in the actomyosin interaction, J. Biol. Chem. 279, 23415–23421.

[54] Doran, M. H., Pavadai, E., Rynkiewicz, M. J., Walklate, J., Bullitt, E., Moore, J. R., Regnier, M., Geeves, M. A., and Lehman, W. (2020) Cryo-EM and Molecular Docking Shows Myosin Loop 4 Contacts Actin and Tropomyosin on Thin Filaments, Biophys. J. 119, 821–830.

