## Supplementary Information for "Natural variants with 2D correlation genetics identify domains coordinating sarcomere proteins during contraction"

Thomas P. Burghardt

Department of Biochemistry and Molecular Biology

200 First St. SW

Mayo Clinic Rochester

Rochester, MN 55905

March 2021

| domain (cd) | seq | index |
|---|---|---|
| M7 actin binding (ab) | <525-563, 647-655> | 1 |
| M7 active site (ac) | <114-125, 167-178, 244-253, 260-267, 453-465, 666-673> | 2 |
| M7 Blocked Head/Converter (bh) | various 295-397 | 3 |
| M7 C-loop (cl) | 359-377 | 4 |
| M7 Converter (cv) | 711-768 | 5 |
| M7 SH3 (h3) | 28-78 | 6 |
| M7 20k (k2) | 634-843 | 7 |
| M7 50k (k5) | 209-633 | 8 |
| M7 27k (k7) | Nterm-208 | 9 |
| M7 Lever-arm (la) | 769-843 | 10 |
| M7 LMM (lm) | 1357-Cterm | 11 |
| M7 Loop 1 (l1) | 202-212 | 12 |
| M7 Loop 2 (l2) | 622-646 | 13 |
| M7 Loop 3 (l3) | 564-576 | 14 |
| M7 MESA (me) | various 168-664 | 15 |
| M7 Myopathy-loop (ml) | 400-414 | 16 |
| M7 MESA-trail (mr) | various 172-917 | 17 |
| M7 S2-C1 mybpc3 (m1) | 844-857 | 18 |
| M7 S2-LT mybpc3 (m2) | 924-936 | 19 |
| M7 S2-C2 mybpc3 (m3) | 858-870 | 20 |
| M7 LMM-Cx (m5) | 1554-1581 | 21 |
| M7 OM binding (om) | various 91-712 | 22 |
| M7 IQ ELC (qe) | 788-798 | 23 |
| M7 IQ RLC (qr) | 814-824 | 24 |
| M7 SH1/SH2 hinge (sh) | 683-710 | 25 |
| M7 Switch 2 helix (sw) | 466-505 | 26 |
| M7 Subfragment 2 (s2) | 844-1356 | 27 |
| M7 LMM-titin (t1) | 1815-1831 | 28 |
| M7 LMM-myomesin (y1) | 1506-1674 | 29 |
| M7 N-term ELC (en) | Nterm-60 | 30 |
| M7 EF1 ELC (e1) | 49-86 | 31 |
| M7 linker ELC (el) | 87-127 | 32 |
| M7 EF2 ELC (e2) | 128-163 | 33 |
| M7 EF3 ELC (e3) | 164-Cterm | 34 |
| M7 N-term RLC (rn) | Nterm-23 | 35 |
| M7 EF1 RLC (r1) | 24-59 | 36 |
| M7 linker RLC (rl) | 60-94 | 37 |
| M7 EF2 RLC (r2) | 95-129 | 38 |
| M7 EF3 RLC (r3) | 130-166 | 39 |
| C3 phospho Ser 1 (s1) | 43-51 | 40 |
| C3 c0-Ig like (c0) | 1-101 | 41 |
| C3 proline rich (pr) | 102-152 | 42 |
| C3 zinc site 1 (z1) | <208, 210, 223, 225> | 43 |
| C3 c1-Ig like (c1) | 153-256 | 44 |
| C3 phospho Ser 2 (st) | 271-279 | 45 |
| C3 phospho Ser 3 (s3) | 280-288 | 46 |
| C3 phospho Ser 4 (s4) | 300-307 | 47 |
| C3 phospho Ser 5 (s5) | 308-315 | 48 |
| C3 linker c1-c2 (lt) | 257-361 | 49 |
| C3 phospho Ser 6 (s6) | 423-431 | 50 |
| C3 c2-Ig like (c2) | 362-452 | 51 |
| C3 c3-Ig like (c3) | 453-543 | 52 |
| C3 phospho Ser 7 (s7) | 546-554 | 53 |
| C3 phospho Thr 8 (s8) | 603-611 | 54 |
| C3 c4-Ig like (c4) | 544-633 | 55 |
| C3 linker c4-c5 (l5) | 634-644 | 56 |
| C3 c5-Ig like (c5) | 645-711 | 57 |
| C3 linker c5-f6 (l6) | 712-743 | 58 |
| C3 c6-Fibronectin (f6) | 744-870 | 59 |
| C3 c7-Fibronectin (f7) | 871-967 | 60 |
| C3 linker c7-c8 (l8) | 968-970 | 61 |
| C3 c8-Ig like (c8) | 971-1065 | 62 |
| C3 c9-Fibronectin (f9) | 1066-1163 | 63 |
| C3 linker c9-c10 (lx) | 1164-1180 | 64 |
| C3 c10-Ig like (cx) | 1181-1274 | 65 |

**Table S1.** Protein domain names, two letter codes (cd), protein sequence assignment (seq), and indexed numbering for domains in the complex βmys/MYBPC3. Domain names begin with M7 or C3 designating origin from βmys or MYBPC3, respectively. Myosin default domains 27k, 50k, and 20k refer to the tryptic proteolytic fragments from cleavage of the MHC sequence in Loop 1 at the active site and Loop 2 in the actin binding site [1]. Most domains are identified within the idealized overall protein structure in **Fig 2**.

| code | population | index |
|---|---|---|
| ACP | ACPOP | 1 |
| AFM | African American | 2 |
| AFR | African | 3 |
| AMR | American | 4 |
| ASI | Asian | 5 |
| ASJ | Ashkenazi Jewish | 6 |
| CEA | CentralAmerican | 7 |
| CUB | Cuban | 8 |
| DOM | Dominican | 9 |
| EAS | East Asian | 10 |
| EST | Estonian | 11 |
| EUA | European American | 12 |
| EUR | European | 13 |
| GLO | Global | 14 |
| MEX | Mexican | 15 |
| NAM | NativeAmerican | 16 |
| NHI | NativeHawaiian | 17 |
| OTH | Other | 18 |
| PCC | PARENT AND CHILD COHORT | 19 |
| PRI | PuertoRican | 20 |
| SAS | SouthAsian | 21 |
| SOA | SouthAmerican | 22 |
| TWC | TWIN COHORT | 23 |

**Table S2**. Human population codes and descriptions from 1000Genomes, gnomAD-Genomes, TopMed, and other studies (see below) in the NCBI database pertaining to βmys and MYBPC3 SNP variants. ACPOP (ACP) whole-genome sequenced control population study from Västerbotten County in northern Sweden; Dominican (DOM) Dominican Republic; Global (GLO) worldwide aggregate missense SNV data from sequencing studies that vary by SNV. Other (OTH) missense SNVs where a population category is not indicated; Parent and Child Cohort (PCC) the UK10K Avon Longitudinal Study of Parents and Children Variants; Twin Cohort (TWC) the UK10K Department of Twin Research and Genetic Epidemiology twin registry of 11,000 identical and non-identical twins between the ages of 16 and 85 years.

| code | new phenotypes | index |
|---|---|---|
| al | Abnormal morphology of left ventricular trabeculae | 1 |
| ar | Arrhythmia | 2 |
| at | Abnormality of T cell physiology | 3 |
| av | Arrhythmogenic right ventricular cardiomyopathy, type 9 | 4 |
| bs | Brugada syndrome | 5 |
| ca | Camptocormia | 6 |
| cc | Catecholaminergic polymorphic ventricular tachycardia type 1 | 7 |
| cm | Cardiomyopathy | 8 |
| cn | Chest pain | 9 |
| co | Congenital myopathy | 10 |
| cp | Cardiovascular phenotype | 11 |
| cr | Cardiac arrest | 12 |
| dc | Primary dilated cardiomyopathy | 13 |
| dg | Delayed gross motor development | 14 |
| dm | Myopathy, distal, 1 (MYH7) | 15 |
| dp | Dyspnea | 16 |
| fd | First degree atrioventricular block | 17 |
| hb | Hyaline body myopathy | 18 |
| hc | Familial hypertrophic cardiomyopathy | 19 |
| ig | Inborn genetic diseases | 20 |
| lv | Left ventricular noncompaction cardiomyopathy | 21 |
| m7 | MYH7-Related Disorders | 22 |
| md | Muscular Diseases | 23 |
| ms | Myosin storage myopathy | 24 |
| mt | MYL2-Related Disorders | 25 |
| my | MYBPC3-Related Disorders | 26 |
| pu | Peripheral neuropathy | 27 |
| px | Paroxysmal atrial fibrillation (MYBPC3) | 28 |
| qt | Prolonged QT interval | 29 |
| rc | Familial isolated restrictive cardiomyopathy | 30 |
| sc | Scapuloperoneal myopathy | 31 |
| sd | Death in infancy | 32 |
| tf | Tetralogy of Fallot | 33 |
| ve | Ventricular extrasystoles | 34 |
| wp | Wolff-Parkinson-White pattern | 35 |
| uk | conflicting-interpretations-of-phenotype | 36 |

**Table S3**. Phenotype (ph) with 2 letter codes. Most phenotypes associate with βmys and MYBPC3 SNV's except when an exclusive SNV location is noted in parenthesis. Conflicting-interpretation-of-phenotype (code uk) implies no consensus phenotype from the database.

| domain (cd) | seq | index |
|---|---|---|
| MT Tyrosine phosphorylation @ Y18 1 (y1) | y1 | 1 |
| MT Tyrosine phosphorylation @ Y29 1 (y2) | y2 | 2 |
| MT N-term insert 1 (n1) | n1 | 3 |
| MT N-term insert 2 (n2) | n2 | 4 |
| MT proline rich 1 (p1) | p1 | 5 |
| MT proline rich 2 (p2) | p2 | 6 |
| MT proline rich 3 (p3) | p3 | 7 |
| MT PHF6* hexapeptide (VQIINK) motif (x1) | | 8 |
| MT PHF6 hexapeptide (VQIVYK) motif (x2) | x2 | 9 |
| MT cys291 PHF (s9) | | 10 |
| MT cys322 PHF (s2) | s2 | 11 |
| MT mtbr 1 (t1) | t1 | 12 |
| MT mtbr 2 (t2) | | 13 |
| MT mtbr 3 (t3) | t3 | 14 |
| MT mtbr 4 (t4) | t4 | 15 |
| MT various (vr) | none from above | 16 |
| C3 phospho Ser 1 (s1) | 43-51 | 17 |
| C3 c0-Ig like (c0) | 1-101 | 18 |
| C3 proline rich (pr) | 102-152 | 19 |
| C3 zinc site 1 (z1) | <208, 210, 223, 225> | 20 |
| C3 c1-Ig like (c1) | 153-256 | 21 |
| C3 phospho Ser 2 (st) | 271-279 | 22 |
| C3 phospho Ser 3 (s3) | 280-288 | 23 |
| C3 phospho Ser 4 (s4) | 300-307 | 24 |
| C3 phospho Ser 5 (s5) | 308-315 | 25 |
| C3 linker c1-c2 (lt) | 257-361 | 26 |
| C3 phospho Ser 6 (s6) | 423-431 | 27 |
| C3 c2-Ig like (c2) | 362-452 | 28 |
| C3 c3-Ig like (c3) | 453-543 | 29 |
| C3 phospho Ser 7 (s7) | 546-554 | 30 |
| C3 phospho Thr 8 (s8) | 603-611 | 31 |
| C3 c4-Ig like (c4) | 544-633 | 32 |
| C3 linker c4-c5 (l5) | 634-644 | 33 |
| C3 c5-Ig like (c5) | 645-711 | 34 |
| C3 linker c5-f6 (l6) | 712-743 | 35 |
| C3 c6-Fibronectin (f6) | 744-870 | 36 |
| C3 c7-Fibronectin (f7) | 871-967 | 37 |
| C3 linker c7-c8 (l8) | 968-970 | 38 |
| C3 c8-Ig like (c8) | 971-1065 | 39 |
| C3 c9-Fibronectin (f9) | 1066-1163 | 40 |
| C3 linker c9-c10 (lx) | 1164-1180 | 41 |
| C3 c10-Ig like (cx) | 1181-1274 | 42 |

**Table S4.** Protein domain names, two letter codes (cd), protein sequence assignment (seq), and indexed numbering for domains in the complex MAPT/MYBPC3. Domain names begin with MT or C3 designating origin from MAPT or MYBPC3, respectively. MAPT sequence is not numerical because each isoform has a different sequence. Acronym mtbr is for MAPT microtubule binding repeat. Various domain (vr) includes every residue not falling into otherwise named domains. It is non-functional and ignored in the analysis tracing co-domains in complex MAPT/MYBPC3. All named domains are shown in **Fig 2**.

| code | population (po) | index |
|---|---|---|
| ACP | ACPOP | 1 |
| AFM | African American | 2 |
| AFR | African | 3 |
| AMR | American | 4 |
| ASI | Asian | 5 |
| ASJ | Ashkenazi Jewish | 6 |
| CAS | Central Asia | 7 |
| CAU | Caucasus | 8 |
| CEA | CentralAmerican | 9 |
| CUB | Cuban | 10 |
| DAG | Daghestan | 11 |
| DAN | Danish | 12 |
| DOM | Dominican | 13 |
| EAS | East Asian | 14 |
| EST | Estonian | 15 |
| EUA | European American | 16 |
| EUR | European | 17 |
| FIN | Finnish from FINRISK project | 18 |
| GLO | Global | 19 |
| GON | Genome of the Netherlands | 20 |
| KOR | KOREAN | 21 |
| LA1 | Latin American 1 | 22 |
| LA2 | Latin American 2 | 23 |
| MDE | Middle_Est | 24 |
| MEX | Mexican | 25 |
| NAM | NativeAmerican | 26 |
| NHI | NativeHawaiian | 27 |
| NRE | Near_East | 28 |
| OCE | Oceania | 29 |
| OTH | Other | 30 |
| PCC | PARENT AND CHILD COHORT | 31 |
| PRI | PuertoRican | 32 |
| SAS | South Asian | 33 |
| SOA | SouthAmerican | 34 |
| SPC | Spanish controls | 35 |
| TWC | TWIN COHORT | 36 |

**Table S5**. Human population codes and descriptions from 1000Genomes, gnomAD-Genomes, TopMed, and other studies (see below) in the NCBI database pertaining to MAPT and MYBPC3

SNP variants. ACPOP (ACP) whole-genome sequenced control population study from Västerbotten County in northern Sweden; Dominican (DOM) Dominican Republic; Global (GLO) worldwide aggregate missense SNV data from sequencing studies that vary by SNV. Other (OTH) missense SNVs where a population category is not indicated; Parent and Child Cohort (PCC) the UK10K Avon Longitudinal Study of Parents and Children Variants; Twin Cohort (TWC) the UK10K Department of Twin Research and Genetic Epidemiology.

| code | new phenotypes | index |
| --- | --- | --- |
| am | Amyloidogenic transthyretin amyloidosis (MAPT) | 1 |
| az | Alzheimer disease (MAPT) | 2 |
| bs | Brugada syndrome | 3 |
| cc | Catecholaminergic polymorphic ventricular tachycardia type 1 | 4 |
| cm | Cardiomyopathy | 5 |
| cp | Cardiovascular phenotype | 6 |
| cr | Cardiac arrest | 7 |
| cs | Conduction disorder of the heart | 8 |
| d1 | Dilated cardiomyopathy 1A (MYBPC3) | 9 |
| dc | Primary dilated cardiomyopathy | 10 |
| ft | Frontotemporal dementia (MAPT) | 11 |
| h1 | Familial hypertrophic cardiomyopathy 1 | 12 |
| h4 | Familial hypertrophic cardiomyopathy 4 (MYBPC3) | 13 |
| hc | Familial hypertrophic cardiomyopathy | 14 |
| hs | Hirschsprung disease | 15 |
| ig | Inborn genetic diseases | 16 |
| lc | Left ventricular noncompaction 10 (MYBPC3) | 17 |
| lv | Left ventricular noncompaction cardiomyopathy | 18 |
| mp | MAPT-Related Spectrum Disorders | 19 |
| my | MYBPC3-Related Disorders | 20 |
| pd | Pick's disease (MAPT) | 21 |
| pl | Parkinson disease, late-onset (MAPT) | 22 |
| pu | Peripheral neuropathy | 23 |
| px | Paroxysmal atrial fibrillation (MYBPC3) | 24 |
| qt | Prolonged QT interval | 25 |
| so | Progressive supranuclear ophthalmoplegia (MAPT) | 26 |
| ss | Shy-Drager syndrome (MAPT) | 27 |
| tp | Malignant tumor of prostate | 28 |
| wp | Wolff-Parkinson-White pattern | 29 |
| uk | conflicting-interpretations-of-phenotype | 30 |

**Table S6**. Phenotype (ph) with 2 letter codes. Phenotypes associated exclusively with MAPT or MYBPC3 are indicated with the gene name in parenthesis. Conflicting-interpretation-of-phenotype (code uk) implies no consensus phenotype from the database.

| Population Sequence | Code | Distance from Addis Ababa km | Line km | Limits km | Outlier | Index |
|---|---|---|---|---|---|---|
| African | AFR | 3763 | 3763 | {1218, 6879} | | 1 |
| Ashkenazi Jewish | ASJ | 4633 | 4633 | {2538, 8006} | | 2 |
| European | EUR | 5550 | 5502 | {5550, 8006} | * | 3 |
| PARENT AND CHILD COHORT | PCC | 6372 | 6372 | {6360, 9241} | | 4 |
| European American | EUA | 7241 | 7241 | {6503, 9453} | | 5 |
| SouthAsian | SAS | 8111 | 8111 | {6067, 9643} | | 6 |
| Asian | ASI | 8981 | 8981 | {6710, 10 565} | | 7 |
| American | AMR | 9850 | 9850 | {6843, 10 639} | | 8 |
| ACPOP | ACP | 10 720 | 10 720 | {7566, 10 956} | | 9 |
| African American | AFM | 11 589 | 11 589 | {5740, 11 980} | | 10 |
| Cuban | CUB | 12 459 | 12 459 | {7839, 12 544} | | 11 |
| Estonian | EST | 13 329 | 13 329 | {8045, 13 980} | | 12 |
| PuertoRican | PRI | 14 198 | 14 198 | {8810, 14 254} | | 13 |
| East Asian | EAS | 14 852 | 15 068 | {9794, 14 852} | * | 14 |
| Dominican | DOM | 15 938 | 15 938 | {10 527, 17 268} | | 15 |
| SouthAmerican | SOA | 16 807 | 16 807 | {11 430, 18 040} | | 16 |
| CentralAmerican | CEA | 17 677 | 17 677 | {13 051, 20 795} | | 17 |
| Mexican | MEX | 18 546 | 18 546 | {14 105, 22 484} | | 18 |
| NativeAmerican | NAM | 19 416 | 19 416 | {16 282, 27 580} | | 19 |
| NativeHawaiian | NHI | 20 404 | 20 286 | {20 404, 30 634} | * | 20 |

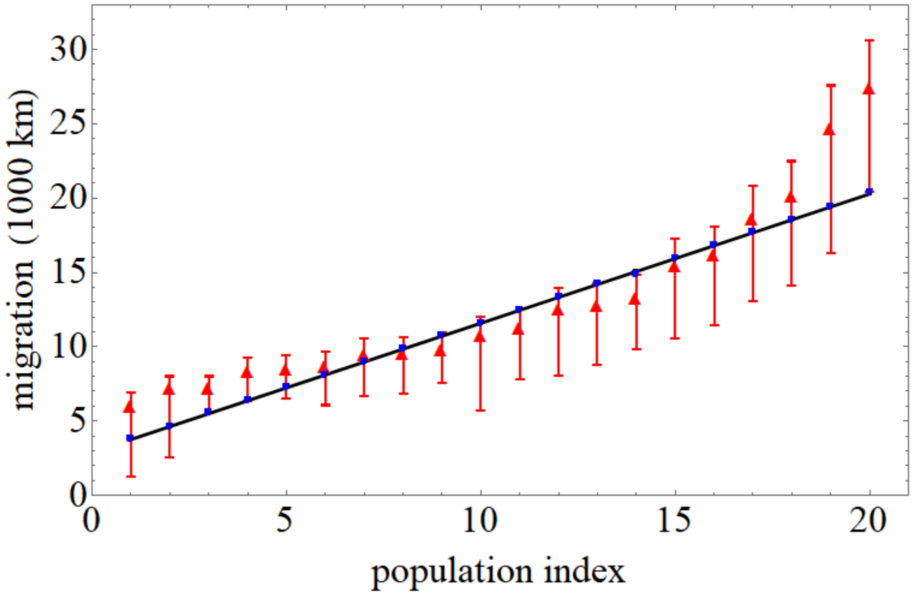


**Table S7**. Migration distance vs human population index for the **Table S2** data set in tabular (top) and graphical (bottom) forms. **Top**. Populations (column 1) listed are in a linear relationship with migration distance (column 3). The migration distance is a proxy for genetic diversity variation where diversity decreases with distance. Fitted line (column 4), computed as described in the text, closely follows migration distances (column 3) and falls within limits (column 5) except for outliers identified with the asterisk in column 6. We use the population index (column 7) to identify populations in graphical presentations like that in the bottom panel. **Bottom**. Blue line indicates a best estimate for the linear relationship of listed populations with migration distance. Red vertical bars at each triangle show minimum limits needed to fulfill the linear estimate as described in the text. Red triangles indicate migration distances falling within the red vertical bars best fitted by the blue line.

**a**

| Population Sequence | Code | Distance from Addis Ababa km | Line km | Limits km | Outlier | Index |
|---|---|---|---|---|---|---|
| Near_East | NRE | 3228 | 2985 | {3228, 5188} | * | 1 |
| Middle_Est | MDE | 3519 | 3519 | {3518, 5936} | | 2 |
| Caucasus | CAU | 4052 | 4052 | {3729, 6480} | | 3 |
| Daghestan | DAG | 4586 | 4586 | {3897, 6642} | | 4 |
| African | AFR | 5119 | 5119 | {1190, 7002} | | 5 |
| Ashkenazi Jewish | ASJ | 5652 | 5652 | {2480, 8149} | | 6 |
| European | EUR | 6186 | 6186 | {5424, 8149} | | 7 |
| Central Asia | CAS | 6719 | 6719 | {4947, 8410} | | 8 |
| Genome of the Netherlands | GON | 7252 | 7252 | {6000, 9039} | | 9 |
| Danish | DAN | 7786 | 7786 | {6122, 9182} | | 10 |
| PARENT AND CHILD COHORT | PCC | 8319 | 8319 | {6215, 9406} | | 11 |
| European American | EUA | 8853 | 8853 | {6355, 9622} | | 12 |
| SouthAsian | SAS | 9386 | 9386 | {5929, 9815} | | 13 |
| Spanish controls | SPC | 9919 | 9919 | {6703, 10013} | | 14 |
| Asian | ASI | 10453 | 10453 | {6558, 10754} | | 15 |
| American | AMR | 10829 | 10986 | {6687, 10829} | * | 16 |
| Oceania | OCE | 11520 | 11520 | {7411, 11531} | | 17 |
| African American | AFM | 12053 | 12053 | {5610, 12193} | | 18 |
| Latin American 1 | LA1 | 12586 | 12586 | {5282, 12662} | | 19 |
| Cuban | CUB | 12768 | 13120 | {7661, 12768} | * | 20 |
| Estonian | EST | 13653 | 13653 | {7862, 14229} | | 21 |
| PuertoRican | PRI | 14186 | 14186 | {8609, 14509} | | 22 |
| Finnish from FINRISK project | FIN | 14640 | 14720 | {7869, 14640} | * | 23 |
| East Asian | EAS | 15118 | 15253 | {9572, 15118} | * | 24 |
| Korean | KOR | 15787 | 15787 | {9548, 16416} | | 25 |
| Dominican | DOM | 16320 | 16320 | {10288, 17577} | | 26 |
| SouthAmerican | SOA | 16853 | 16853 | {11171, 18362} | | 27 |
| Latin American 2 | LA2 | 17387 | 17387 | {12425, 19697} | | 28 |
| CentralAmerican | CEA | 17920 | 17920 | {12755, 21166} | | 29 |
| Mexican | MEX | 18453 | 18453 | {13784, 22886} | | 30 |
| NativeAmerican | NAM | 18987 | 18987 | {15912, 28073} | | 31 |
| NativeHawaiian | NHI | 19940 | 19520 | {19940, 31181} | * | 32 |

**Table S8.** Migration distance vs human population index for the **SI Table S5** data set in tabular (**panel a**) and graphical (**panel b**) forms. **Panel a**. Populations (column 1) listed are in a linear relationship with migration distance (column 3). The migration distance is a proxy for genetic diversity variation where diversity decreases with distance. Fitted line (column 4), computed as described in the text, closely follows migration distances (column 3) and falls within limits (column 5) except for outliers identified with the asterisk in column 6. We use the population index (column 7) to identify populations in graphical presentations like that in **panel b** below.

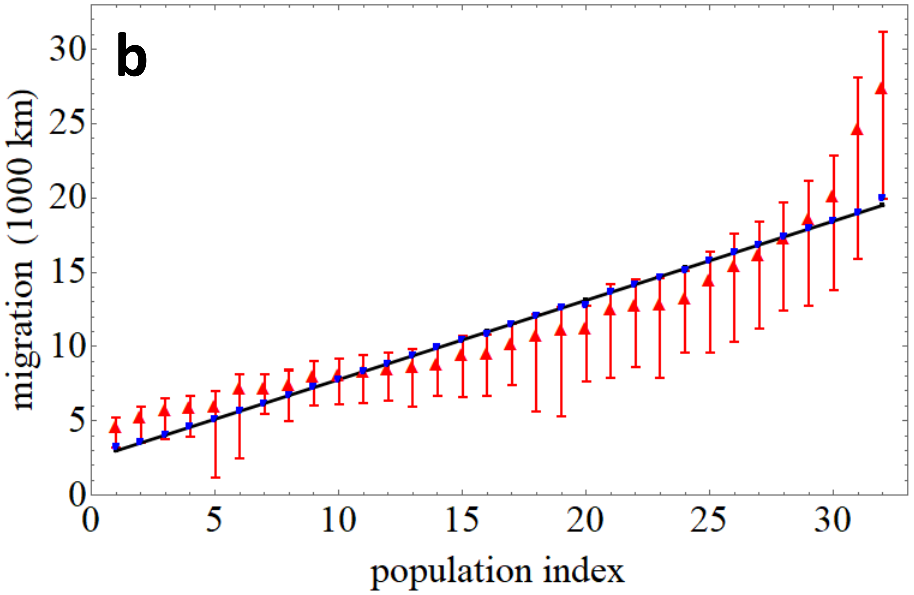


**Panel b**. Blue line indicates a best estimate for the linear relationship of listed populations with migration distance. Red vertical bars at each triangle show minimum limits needed to fulfill the linear estimate as described in the text. Red triangles indicate migration distances falling within the red vertical bars best fitted by the blue line.

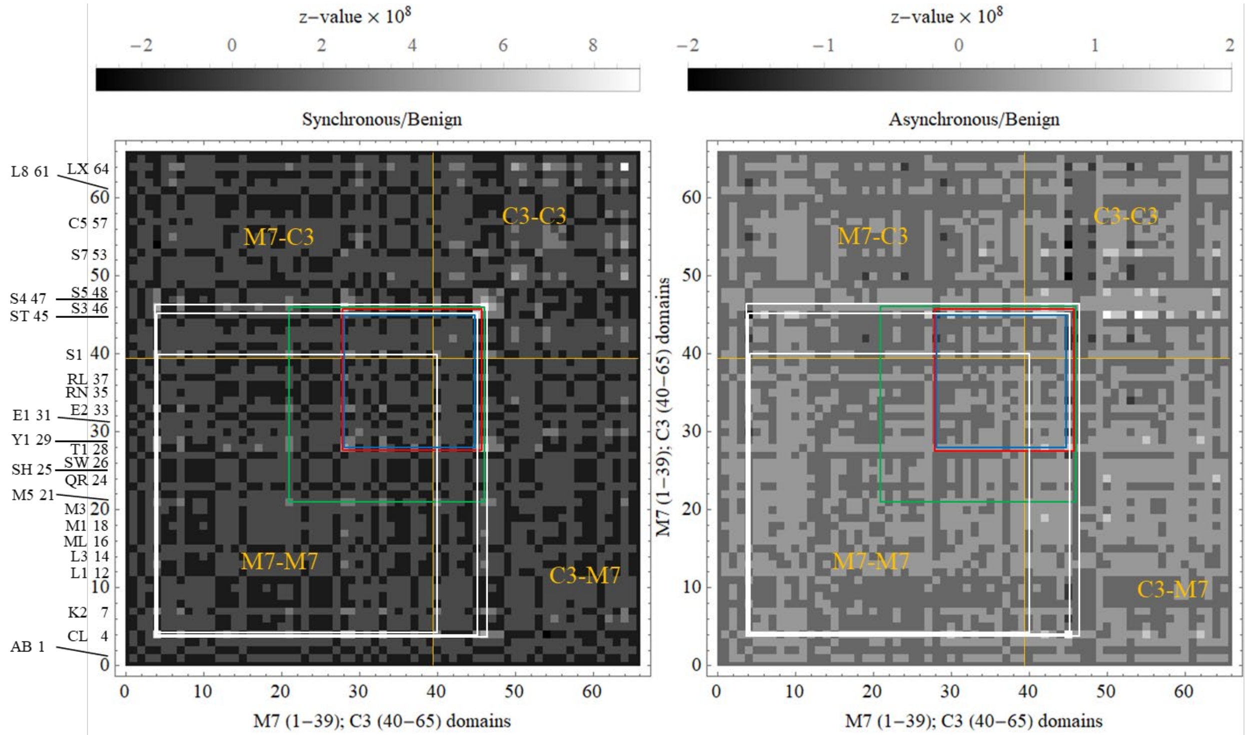


**Figure S1.** 2D correlation maps for complex βmys/MYBPC3 with benign outcomes. Axes represent βmys (M7 1-39) followed by MYBPC3 (C3 40-65) domains. M7 and C3 abbreviate βmys and MYBPC3, respectively. Domain name, two letter code, protein sequence, and index number are from **SI Table S1** and several protein domains are also labeled by their two-letter code on the leftmost axis in the figure. Intensities (z-values) are indicated numerically by the grayscale. The 4 regions defined by vertical and horizontal orange lines at the interface of pixels 39-40 labeled M7-M7, M7-C3, C3-C3, and C3-M7 separate inter-protein cross-peaks (M7-M7 and C3-C3) from intra-protein cross-peaks (M7-C3 and C3-M7). The 6 most significant pathogenic co-domains are linked by correlation squares (white, green, red, blue, for different βmys domains then repeating color sequence as needed) that link the off-diagonal interacting domain coordinates falling within M7-C3 and C3-M7 regions.

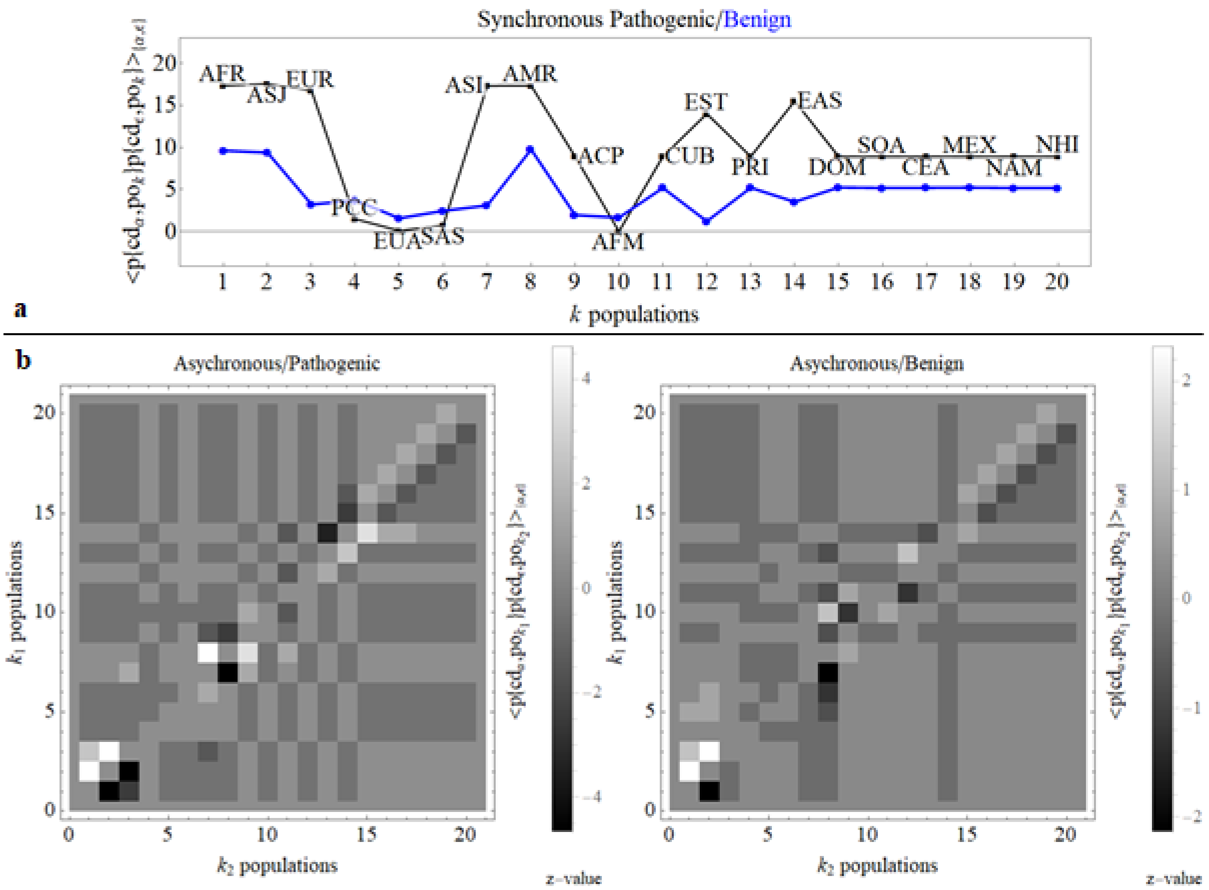


**Figure S2.** Population dependence for co-domains using cross-peak intensity averaged over the 6 most significant pathways identified by pathogenic (**Fig 7 panel a**) or benign (**Fig 8 panel a**) SNV's. **Panel a** has synchronous interactions for pathogenic (black) or benign (blue) SNV's. **Panel b** has asynchronous interactions for pathogenic (left) or benign (right) SNV's. Population three letter code and index are indicated in **panel a**.

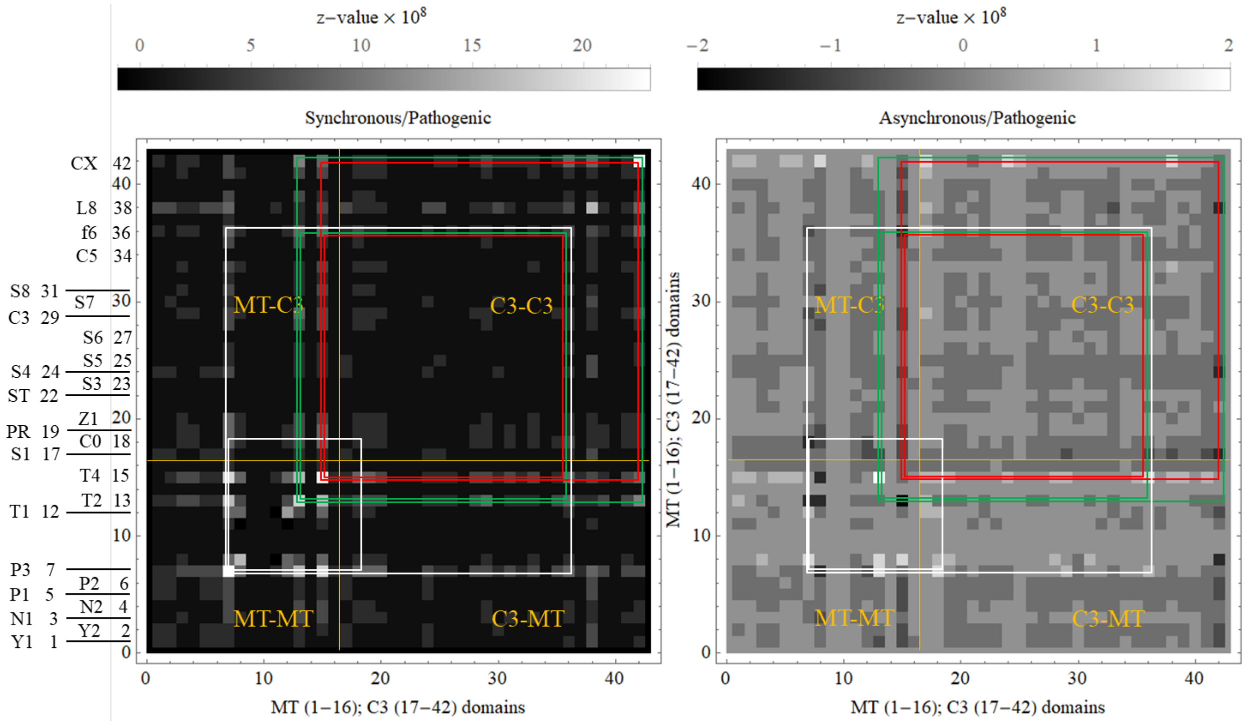


**Figure S3.** 2D correlation maps for complex MAPT/MYBPC3 with pathogenic outcomes. Axes represent MAPT (MT 1-16) followed by MYBPC3 (C3 17-42) domains where MT and C3 abbreviates MAPT and MYBPC3, respectively. Domain name, two letter code, protein sequence, and index number are from **SI Table S4** and several protein domains are also labeled by their two-letter code on the leftmost axis in the figure. Intensities (z-values) are indicated numerically by the grayscale. The 4 regions defined by vertical and horizontal orange lines at the interface of pixels 39-40 labeled MT-MT, MT-C3, C3-C3, and C3-MT separate inter-protein cross-peaks (MT-MT and C3-C3) from intra-protein cross-peaks (MT-C3 and C3-MT). The 6 most significant pathogenic co-domains are linked by correlation squares (white, green, red, blue, for different MAPT domains then repeating color sequence as needed) that link the off-diagonal interacting domain coordinates falling within MT-C3 and C3-MT regions.

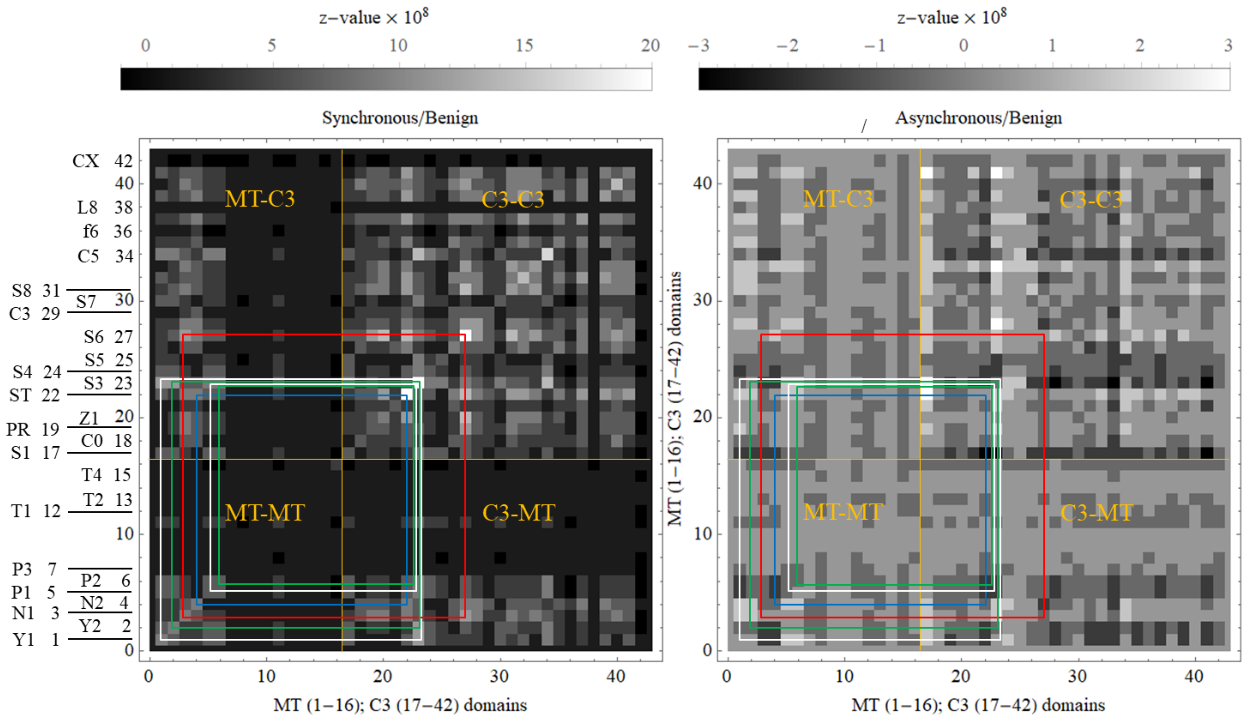


**Figure S4.** 2D correlation maps for complex MAPT/MYBPC3 with benign outcomes. Nomenclature is otherwise identical to **Fig S3**.

**Data Sets**

1. Name: 6ddp complex βmys/MYPBC3

Caption: Fulfilled and unknown 6 dimensional data points (6ddp) for complex βmys/MYPBC3

Description: Fulfilled and unknown 6 dimensional data points (6ddp) for complex βmys/MYPBC3 containing mutation site, residue substitution, phenotype, and pathogenicity.

File name: 6ddpbmysMYBPC3.xls

2. Name:6ddp complex MAPT/MYBPC3

Caption: Fulfilled and unknown 6 dimensional data points (6ddp) for complex MAPT/MYBPC3

Description: Fulfilled and unknown 6 dimensional data points (6ddp) for complex MAPT/MYBPC3 containing mutation site, residue substitution, phenotype, and pathogenicity.

File name: 6ddpMAPTMYBPC3.xls

Reference

[1] Balint, M., Sreter, F. A., Wolf, I., Nagy, B., and Gergely, J. (1975) The substructure of heavy meromyosin. The effect of Ca 2+ and Mg 2+ on the tryptic fragmentation of heavy meromyosin, *J. Biol. Chem. 250*, 6168-6177.
